# A likelihood ratio test for detecting shifts in homeolog expression ratios in allopolyploids

**DOI:** 10.1101/2025.07.01.660977

**Authors:** Jianqiang Sun, Jun Sese, Kentaro K. Shimizu

## Abstract

Allopolyploids arise through hybridization between related species, carrying multiple sets of chromosomes from distinct progenitor, referred to as subgenomes. Within allopolyploids, duplicated genes across subgenomes, called homeologs, are thought to enhance environmental robustness by shifting their expression ratios depending on environmental and developmental changes. However, existing methods for detecting such ratio shifts, including HomeoRoq and Fisher’s exact test, are limited to allopolyploids with two subgenome sets inherited from two progenitors, and thus cannot handle more complex cases (e.g., allohexaploid wheat). Here, we present the HOmeolog Bias Identification Test (HOBIT), a statistical method for detecting shifts in homeolog expression ratios across different conditions using RNA-Seq count data. HOBIT performs a likelihood ratio test for each homeolog, comparing a full model allowing homeolog expression ratios to vary across conditions with a reduced model assuming constant ratios. Simulation benchmarks for allotetraploids and allohexaploids demonstrated that HOBIT outperforms existing methods in both area under the receiver operating characteristic curve and F1 score. Application to real RNA-Seq datasets from allotetraploid *Cardamine flexuosa*, allotriploid *Cardamine insueta*, and *Triticum aestivum* (wheat) produced biologically consistent results reflecting experimental settings. HOBIT provides a promising framework for uncovering homeolog regulation and adaptive responses in allopolyploids, without restrictions of ploidy complexities.

## Introduction

Allopolyploids play a critical role in global agriculture, including major crops such as wheat, cotton, and canola, which support human populations worldwide as well as in natural species (Adams & Wendel, 2005; Soltis *et al*., 2015; Shimizu, 2022; Akagi *et al*., 2022; Deb *et al*., 2023). They arise through the hybridization of two or more closely related species, resulting in multiple sets of chromosomes derived from distinct progenitor genomes, referred to as subgenomes. Within allopolyploids, duplicated gene copies originating from distinct subgenomes, known as homeologs, can obtain diverse evolutionary trajectories and functions (Adams & Wendel, 2005; Soltis *et al*., 2015; Deb *et al*., 2023). While some homeologs retain balanced homeolog expression ratios (HERs) across subgenomes, others display shifts in HERs depending on tissue type, developmental stage, habitat, and environmental changes. Such biased regulation of HERs is thought to undergo dynamic changes or accumulate changes over generations, potentially increasing the adaptive capacity of allopolyploids and enabling them to thrive across a broader range of environmental conditions than their progenitor species (Shimizu, 2022; Tossi *et al*., 2022).

Studies of homeologs exhibiting shifts in HERs across subgenomes under varying conditions, referred to as ratio-shifted homeologs (RSHs), have motivated the development of statistical methods for their detection. While many methods have been developed to quantify homeolog expression (Page *et al*., 2013; Akama *et al*., 2014; Boatwright *et al*., 2018; Kuo *et al*., 2020; Hu *et al*., 2021; Sun *et al*., 2023), few statistical approaches specifically address RSH detection. One such method is provided by HomeoRoq, which estimates the probability that HERs remain unchanged between two conditions using RNA-Seq read count data for each homeolog (Akama *et al*., 2014). In RNA-Seq studies, gene and homeolog expression are typically quantified by counting sequence reads mapped to specific genomic regions. These count data often exhibit overdispersion, where the variance exceeds the mean. To account for this, HomeoRoq assumes homeolog expression as following a hierarchical normal distribution, with the variance modelled as a function of the mean. HomeoRoq has been applied to various allotetraploid species, including those in the genus *Arabidopsis* and its related genus *Cardamine* (Akama *et al*., 2014; Paape *et al*., 2016; Akiyama *et al*., 2021).

*Cardamine flexuosa* (2*n* = 4*x* = 32, HHAA) is an allotetraploid species that arose from hybridization between *Cardamine amara* (2*n* = 2*x* = 16, AA), which inhabits in wet environments such as running water, and *Cardamine hirsuta* (2*n* = 2*x* = 16, HH), which is adapted to dry habitats such as roadsides (Mandáková *et al*., 2014; Shimizu-Inatsugi *et al*., 2017). Unlike its progenitors, *C. flexuosa* occupies a broader range of environments. The application of HomeoRoq to detect RSHs in *C. flexuosa* across wet and dry habitats revealed that a small proportion of homeologs (0.2%–3.3%) exhibited shifts in HERs between the two habitats (Akiyama *et al*., 2021). Specifically, the RSHs were associated with responses to water deprivation, oxidative stress, reactive oxygen species, and abscisic acid.

Meanwhile, *Cardamine insueta* (2*n* = 3*x* = 24, RRA) has an uneven subgenome composition. One subgenome originated from *C. amara* (A-subgenome), whereas the other two were derived from *Cardamine rivularis* (2*n* = 2*x* = 16, RR) (R-subgenomes), a species that inhabits grassy areas away from riversides (Urbanska & Landolt, 1970; Urbanska-Worytkiewicz & Landolt, 1974). Experimental submergence of *C. insueta* was found to shift HERs in some homeologs in response to submergence stress (Sun *et al*., 2020). However, no statistical assessment was performed in the original study due to the absence of suitable methods, as *C. insueta* has an unbalanced number of subgenomes and the dataset lacks biological replicates.

Allohexaploid wheat, *Triticum aestivum* (2*n* = 6*x* = 42; AABBDD), one of the most important food grains worldwide, serves as a valuable model for studying allopolyploids. It originated through hybridization among three species: *Triticum urartu* (2*n* = 2*x* = 14; AA), an unknown species related to *Aegilops speltoides* (2*n* = 2*x* = 14; SS, which is thought to be the progenitor of BB), and *Aegilops tauschii* (2*n* = 2*x* = 14; DD) (Dubcovsky & Dvorak, 2007; Zhao *et al*., 2023; Okada & Shimizu, 2024). Despite recent advances in wheat genome sequencing and annotation (Zimin *et al*., 2017; The International Wheat Genome Sequencing Consortium (IWGSC) *et al*., 2018; Walkowiak *et al*., 2020; Shimizu *et al*., 2021; Zhu *et al*., 2021; Nomura *et al*., 2025), RNA-Seq studies have primarily focused on overall gene or homeolog expression rather than HERs. A primary limitation of this approach is the lack of a robust statistical framework for detecting RSHs in allopolyploids with more than two subgenomes. For instance, current methods, such as HomeoRoq (Akama *et al*., 2014) or Fisher’s exact test (FET), are restricted to binary comparisons.

Despite the growing interest in HER dynamics, broadly applicable statistical methods for detecting homeologs that exhibit shifts across distinct conditions (i.e., RSHs) remain limited. To address this gap, we present the HOmeolog Bias Identification Test (HOBIT), a statistical method designed to detect RSHs in allopolyploids without constrains on subgenome composition. By providing a flexible and statistically rigorous framework for RSH detection, HOBIT is expected to advance our understanding of the molecular mechanisms underlying environmental adaptation in allopolyploids.

## Materials and Methods

### Algorithm of HOBIT

Since RNA-Seq read count data often exhibit overdispersion, modelling these counts using negative binomial (NB) distributions is standard practice (Robinson & Smyth, 2007; Love *et al*., 2014; Chen *et al*., 2025). Accordingly, HOBIT assumes that homeolog expression follows a NB distribution:

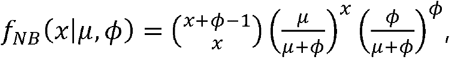

where *μ* and *ϕ* represent the mean and dispersion parameters, respectively.

HOBIT performs the following steps for each homeolog to compute the statistical significance (i.e., *p*-value). 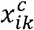 denotes a normalized count representing the homeolog expression of subgenome *k* in biological replicate *i* under condition *c*. For instance, in an experiment with three biological replicates of allohexaploid wheat under control (C) and treatment (T) conditions, the indices are defined as *k* = A,B,D;, *i* = 1,2,3; and *c*= *C,T*. Then, given the mean (*μ* ^*c*^) of gene expression (i.e. the total counts of an homeolog across all subgenomes), dispersion of homeolog expression (*ϕ* ^*c*^), and HER (***θ***^*c*^) under a condition *c*, the likelihood for the observed counts is computed as:

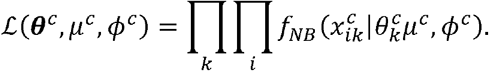

The parameter *μ*^*c*^ is estimated via Markov chain Monte Carlo (MCMC) sampling from non-informative prior (NIP) distributions. The dispersion parameter *ϕ*^*c*^ is estimated using edgeR (Chen *et al*., 2025), which leverages shared information across all homeologs. ***θ***^*c*^ is estimated via MCMC sampling from NIP distributions with the constraint 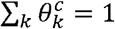.

Next, to statistically determine the RSHs across conditions, full and reduced models are considered. The full model assumes that the HER differs across all conditions. The likelihood is calculated as follows:

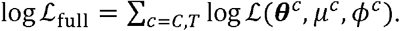

The reduced model, which represents the null hypothesis, is hypothesized to share the same HER under all conditions. The likelihood is calculated as follows:

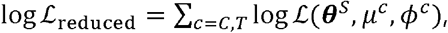

where ***θ***^*S*^ is the average of ***θ***^*c*^ and ***θ***^*T*^. Under the null hypothesis, if HER remains unchanged, then theoretically, ***θ***^*C*^ = ***θ***^*T*^ = ***θ***^*S*^, leading to identical likelihoods for both conditions.

Once the likelihoods are calculated from the full and reduced models, a likelihood ratio test (LRT) is performed to assess the statistical significance:

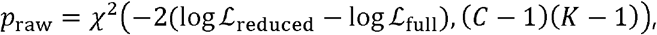

where *C* and *K* denote the number of conditions and subgenomes, respectively. Hereinafter, the resulting test statistic (i.e., *p*_raw_) is referred to as the *p*_raw_-value. As the LRT is known to become increasingly sensitive as sample size increases (Wilks, 1938; Dziak *et al*., 2014; Gudicha *et al*., 2017), even small differences between competing models may yield small *p*_raw_-values. While this can indicate statistically significant differences, such results may not necessarily carry biological meaning. To address this issue, an ad hoc normalization is applied to the log-likelihood difference using a logarithmic function of the number of replicates, as follows:

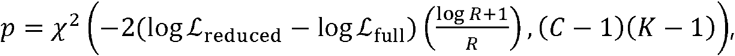

where *R* denotes the average of number of biological replicates per condition. Hereinafter, the resulting test statistic computed using normalized log-likelihoods (i.e., *p*) is referred to as the *p*-value. After processing all homeologs, the *p*-values are adjusted using the Benjamini–Hochberg method, referred to as *q*-values, to control the false discovery rates (Benjamini & Hochberg, 1995).

Furthermore, to quantify the magnitude of HER shifts across conditions for each homeolog, HOBIT also reported absolute differences in HER between conditions, *D*_max_ and *OR*_max_. *D*_max_ is defined as the maximum of absolute difference, representing the largest HER shift observed in at least one subgenome. *OR*_max_ is defined as the maximum odds ratio among any pair of conditions in at least one subgenome.

### HOBIT variants

In addition to the default settings, three alternative configurations of HOBIT are also implemented. First, instead of modelling homeolog expression using the NB distribution, a zero-inflated negative binomial (ZINB) distribution is implemented (Method S1). Second, while the default setting samples 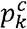 from a NIP distribution under the constraint 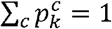, an alternative approach using a Dirichlet prior distribution is also implemented (Method S2). Third, *ϕ*^*c*^ is precomputed using edgeR in the default implementation. As an alternative, *ϕ*^*c*^ is drawn from a NIP distribution.

### Algorithm to simulate artificial benchmark datasets

A series of computational steps was implemented to simulate artificial benchmark datasets. The process began with the preparation of a population of mean and variance values. These values were estimated from real RNA-Seq data: *C. flexuosa* (Akiyama *et al*., 2021) for simulating allotetraploid datasets and wheat (Yang *et al*., 2021) for allohexaploid datasets. Additionally, a quadratic regression model, *f*_*V*_(*μ*), was constructed from the population, in order to model log-transformed variance as a function of the log-transformed mean. From this population, a specified number (*N*) of mean values were randomly sampled to represent baseline expression levels for control and treatment conditions. Next, to introduce differential expression, *N* random fold-change values were drawn from a 1 + gamma distribution and applied randomly to either the control or treatment condition. Then, to simulate RSHs, HERs for the control condition were randomly sampled from a truncated normal distribution, ranging between 0 and 1, with a peak at half for allotetraploid datasets and at one-third for allohexaploid datasets. HERs for the treatment condition were then generated by applying shifts to the control HERs, where the magnitude of shifts was sampled from a hierarchical gamma distribution. Finally, count data were sampled from NB distributions, with means calculated as the product of the adjusted means and HERs, and variances derived from *f*_*V*_ (*μ*) . Ground-truth RSHs were defined based on the criteria *D*_max_ >0.2 or *OR*_max_ >2.0, where *D*_max_ and *OR*_max_ were calculated using the true means. A detailed description of the dataset generation procedure is provided in Method S3 and Figure **S1**.

Using this procedure, 50 artificial RNA-Seq datasets were generated for both allotetraploid species, comprising two subgenomes, and allohexaploid species, comprising three subgenomes. Simulations were conducted under the following conditions: (i) 10,000 homeologs; (ii) two experimental conditions (control and treatment); and (iii) biological replicate counts of three, five, and ten per condition.

### Performance evaluation with artificial datasets

Artificial datasets were analysed using HOBIT and its variants, as well as HomeoRoq and FET, for RSH detection. For HOBIT and its variants, MCMC sampling was conducted using four chains, each with 1,000 warm-up iterations followed by 1,000 sampling iterations. The detection performance was evaluated using two key metrics: the area under the receiver operating characteristic curve (AUC) and F1 score. The AUC was computed as 1 minus the *p*-values, measuring the overall ability of the model to distinguish RSHs from non-RSHs. The F1 score was calculated as the harmonic mean of precision and recall, where precision was defined as the proportion of correctly predicted RSHs among all predicted RSHs, and recall represented the proportion of correctly predicted RSHs among all ground-truth RSHs. The predicted RSHs were defined as homeologs with a *q*-value less than 0.05.

Because HomeoRoq and FET are not designed to handle allohexaploid species with three subgenomes, a one-vs-rest strategy was employed to detect HER shifts in the simulated allohexaploid datasets. Specifically, comparisons were conducted as follows: *θ*_*A*_ vs (*θ*_*B*_ + *θ*_*D*_), *θ* _*B*_ vs (*θ*_*D*_ + *θ*_*A*_), and *θ*_*D*_ vs (*θ*_*A*_ + *θ*_*B*_). The minimum *p*-value among these comparisons was used as the representative *p*-value for each homeolog, and *q*-values were calculated from these *p*-values.

### Case study: Allotetraploid Cardamine flexuosa

To evaluate the performance of HOBIT on real RNA-Seq data, an RNA-Seq study of allotetraploid *C. flexuosa* was selected as a case study. In the original study (Akiyama *et al*., 2021), RNA-Seq was conducted using the leaves of *C. flexuosa* sampled from three distinct ecological habitats: a dry, wet, and intermediate habitats. Sampling was performed on three different dates (April 18, May 2, and May 16 in 2013). Each sample comprises three biological replicates, except for the dry habitat on April 18, where only two biological replicates were available. Homeolog expression was quantified by following HomeoRoq alignment and a classification protocol (Akama *et al*., 2014) from RNA-Seq data and was then normalized to fragments per kilobase of exons per million mapped reads (FPKM). The RSHs between any two habitats for each sampling date were identified by applying HomeoRoq to the FPKM.

In this study, HOBIT was applied to the counts normalized by TMM (Robinson & Oshlack, 2010) and FPKM to detect RSHs between dry and wet habitats. Homeologs with counts less than 1.0 counts per million (CPM) across all libraries were excluded from the analysis. MCMC sampling in HOBIT was performed using ten chains, each with 1,000 warm-up iterations followed by 1,000 sampling iterations. RSHs were defined as homeologs with a *q*-value of <0.05. Additionally, gene ontology (GO) enrichment analysis was conducted using clusterProfiler (Wu *et al*., 2021) against RSHs detected by HOBIT and HomeoRoq using a threshold *p*-value of <0.05. The differences between the RSH sets detected by HOBIT and HomeoRoq were then compared using the Jaccard index and overlap coefficient. The Jaccard index measures the ratio of the intersection size to the union size of the two sets, whereas the overlap coefficient measures the ratio of the intersection size to the size of the smaller set.

The differences between the RSH sets identified by HOBIT and HomeoRoq were then compared using the Jaccard index and the overlap coefficient, where the Jaccard index measures the ratio of the size of the intersection to the size of the union of the two sets, and the overlap coefficient measures the ratio of the size of the intersection to the size of the smaller set.

### Case study: allotriploid Cardamine insueta

To further assess the applicability of HOBIT to allopolyploids with an uneven subgenome composition, we conducted a case study using RNA-Seq data from a previous analysis of the allotriploid *C. insueta* (Sun *et al*., 2020). The original study examined homeolog expression in leaflets from a single biological replicate across nine time points (0, 2, 4, 8, 12, 24, 48, 72, and 96 h) after floating it on water. Statistical detection of RSH was not performed in the original study due to the absence of biological replication.

In this study, read counts of homeolog expression were obtained from a previous study and normalized with TMM (Robinson & Oshlack, 2010). Next, homeologs with a CPM of <1.0 across all replicates were removed.

HOBIT was then applied to identify RSHs across time points with a statistical significance threshold of *p*-value of <0.05. In this context, RSH represents a homeolog that exhibits a shift in its expression ratios at least at one of the nine time points, analogous to analysis of variance (ANOVA). MCMC sampling in HOBIT was conducted using ten chains, each with 1,000 warm-up iterations followed by 1,000 sampling iterations. Additionally, GO enrichment analysis was conducted using clusterProfiler (Wu *et al*., 2021), and the enriched GO in RSHs were defined as GO with a *p*-value of <0.05.

### Case study: Allohexaploid wheat

To demonstrate the applicability of HOBIT to allopolyploids with more than two subgenomes, we further selected a published transcriptomic dataset from allohexaploid wheat that investigated the gene expression associated with heading date variation (Yang *et al*., 2021). In the original study, RNA-Seq was conducted on shoot apex and leaf tissues collected from two homozygous wheat lines that exhibited early and late heading-date phenotypes. Samples were collected at two key developmental stages: W2.0 (double ridge stage) and W3.5 (androgynous primordium differentiation stage), with three biological replicates per sample.

We focused on RNA-Seq data derived from the early heading-date phenotype. Raw reads were pre-processed using Trimmomatic (Bolger *et al*., 2014) with the parameters ‘HEADCROP:14 LEADING:30 TRAILING:30 SLIDINGWINDOW:5:30 MINLEN:60’. The cleaned reads were quantified using the EAGLE-RC pipeline (Kuo *et al*., 2020), which aligns reads to IWGSC RefSeq v1.1 (IWGSC, 2018) using STAR (Dobin *et al*., 2012), followed by subgenome sorting using EAGLE (Kuo *et al*., 2018). The expression values were derived as read counts per homeolog using featureCounts (Liao *et al*., 2014) and normalized with TMM (Robinson & Oshlack, 2010). Homeologs with expression levels below 1.0 CPM across all replicates were excluded from further analysis. HOBIT was then applied to detect RSHs between the two tissues (shoot apex vs. leaf) at each developmental stage, and between the two developmental stages (W2.0 vs. W3.5) within each tissue. MCMC sampling in HOBIT was conducted using ten chains, each with 1,000 warm-up iterations followed by 1,000 sampling iterations. RSHs were defined using a significance threshold of *q*-value <0.05.

### Software tools and execution environment

HOBIT was implemented with Stan v2.36.0 (Stan Development Team, 2024), a Bayesian statistical modelling language and platform. The interface was developed in R v4.5.0 (R Core Team, 2024), a statistical computing and graphics language as well as platform. The source code for both HOBIT and HomeoRoq is available as an R package, *hespresso*, hosted on GitHub https://github.com/bitdessin/hespresso. edgeR v4.6.2 (Chen *et al*., 2025) was internally utilised by HOBIT for dispersion estimation. Additional tools used in case studies of real RNA-Seq data include Trimmomatic v0.39 (Bolger *et al*., 2014), STAR v2.7.10a (Dobin *et al*., 2012), EAGLE v1.1.3 (Kuo *et al*., 2018), EAGLEs-RC v1.1.1 (Kuo *et al*., 2020), featureCounts v2.0.3 (Liao *et al*., 2014), and clusterProfiler v4.16.0 (Wu *et al*., 2021). All analyses were performed using eight parallel threads on Ubuntu 24.04 with an Intel Core i9-11900K processor (8 cores, 16 threads, 3.5 GHz).

## Results

### Performance evaluation with artificial data of allotetraploids

The evaluation for allotetraploid simulated datasets demonstrated that HOBIT consistently achieved the highest AUC, while FET showed the weakest performance (Fig. **1a**). Although the AUCs of all methods improved with increasing numbers of replicates, HOBIT outperformed both HomeoRoq and FET across all replicate conditions. This superior AUC indicates that HOBIT more effectively assigned lower *p*-values to ground-truth RSHs, thereby ranking them higher based on statistical significance.

**Fig. 1.**
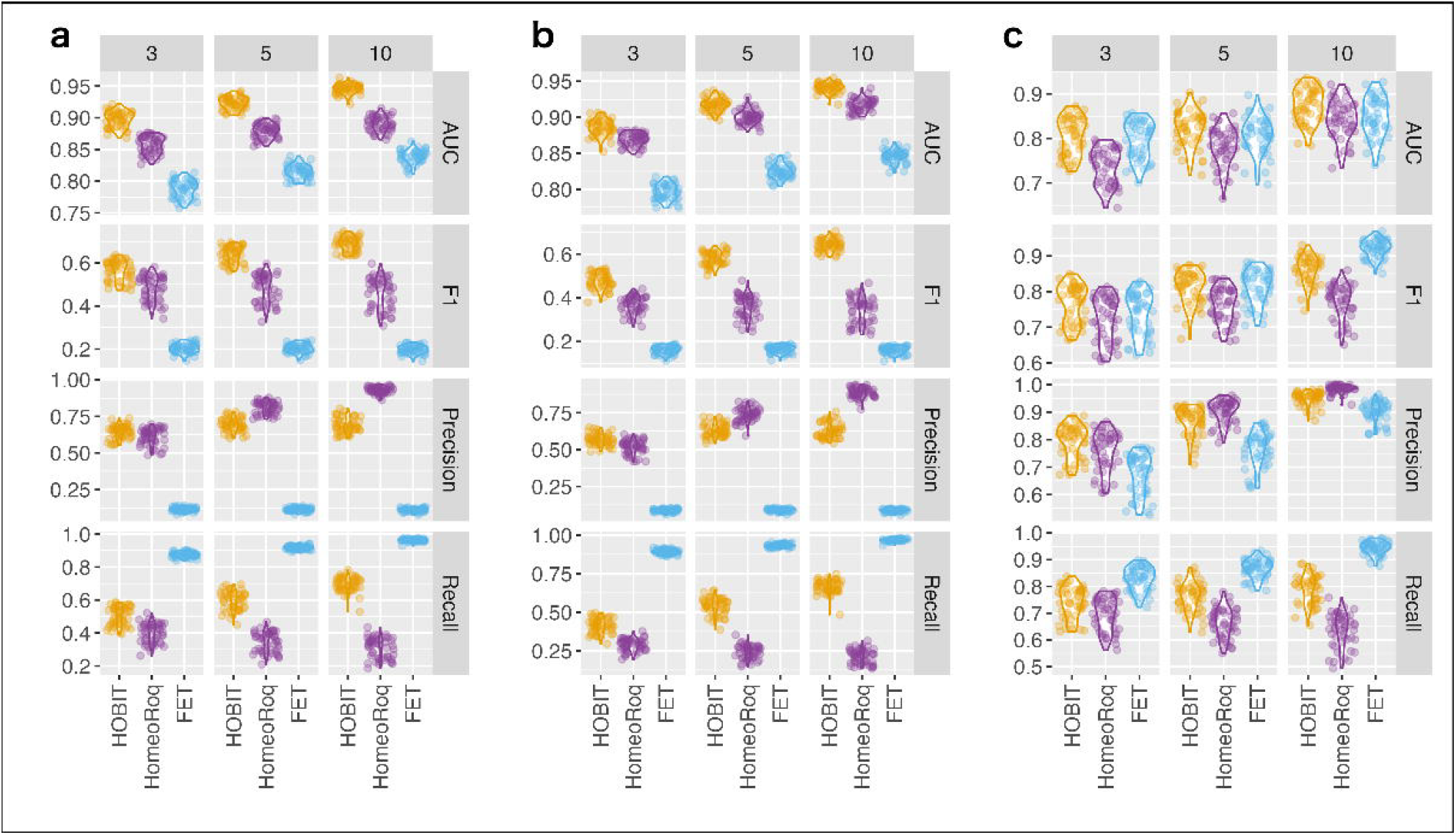
Performance metrics based on 50 artificial datasets simulating an allotetraploid species. Metrics were calculated from (**a**) all homeologs, (**b**) homeologs expressed across all subgenomes and conditions, and (**c**) homeologs with at least one subgenome unexpressed under one of the conditions.

Regarding the F1 score, HOBIT outperformed the other methods. As the number of replicates increased, the F1 score of HOBIT improved steadily, while those of HomeoRoq and FET remained unchanged. As a result, HOBIT exhibited a clear performance advantage at higher replicate counts (e.g., 5 and 10). The robustness of HOBIT stems from its well-balanced trade-off between precision and recall. In contrast, HomeoRoq and FET exhibited imbalanced performance profiles. Specifically, HomeoRoq showed high precision but suffered from low recall, whereas FET showed the opposite trend. With increasing replicate numbers, HomeoRoq improved precision further but at the cost of reduced recall, while FET gained recall with declining precision. These opposing trends limited F1 score improvements for both methods.

Additionally, we found that incorporating a *q*-value threshold alongside an additional cutoff (i.e. *D*_max_ >0.2 or *OR*_max_ >2) in HOBIT improved the F1 score by enhancing precision while retaining high recall, especially with a larger number of replicates (HOBIT+ in Fig. **S2a**). Incidentally, applying the same cutoff to HomeoRoq did not lead to improved F1 score (HomeoRoq+ in Fig. **S2a**).

When restricting the analysis to homeologs expressed across all subgenomes under both conditions, performance trends for AUC, F1 score, precision, and recall were consistent with those observed in the complete dataset (Fig. **1b**). In this subset as well, the application of an additional cutoff improved the F1 score of HOBIT (HOBIT+ in Fig. **S2b**). In contrast, in scenarios where one or more homeologs were not expressed in either condition, HOBIT and FET showed similar performance in terms of both AUC and F1 score, whereas HomeoRoq exhibited lower performance for both metrics, with one notable exception (Fig. **1c**). Specifically, FET achieved the highest F1 score when ten replicates were available. This is because increasing the number of replicates amplifies the total counts for expressed versus unexpressed homeologs, which enhances the bias in the contingency table and consequently results in FET producing lower *p*-values. In this scenario, applying the additional cutoff did not improve the performance of either HOBIT or HomeoRoq (HOBIT+ and HomeoRoq+ in Fig. **S2c**).

Using different definitions resulted in varying sets of ground-truth RSHs with differing magnitudes of HER shift. To evaluate whether detection performance was influenced by the thresholds used to define ground-truth RSHs (*D*_max_ >0.2 or *OR*_max_ >2.0), additional validations were performed using alternative criteria: *D*_max_ >0.2 or *OR*_max_ >1.5, and *D*_max_ >0.2 or *OR*_max_ >3.0. As a result, HOBIT consistently outperformed the other methods in terms of AUC, regardless of the threshold settings. In terms of F1 score, HOBIT outperformed HomeoRoq under *D*_max_ >0.2 or *OR*_max_ >1.5 threshold (Fig. **S3**) and showed comparable performance under *D*_max_ >0.2 or *OR*_max_ >3.0 thresholds (Fig. **S4**), while FET consistently showed the lowest performance across all threshold settings. Notably, applying more stringent additional cutoff aligned with the simulation criteria (e.g., *D*_max_ >0.2 or *OR*_max_ >3.0) improved the F1 score of HOBIT when the number of replicates was large (HOBIT+ in Fig. **S5**).

### Performance evaluation with artificial data of allohexaploids

The overall validation trends observed in artificial datasets simulating allohexaploid species were consistent with those from allotetraploid simulations (Fig. **2**). In particular, HOBIT consistently outperformed HomeoRoq in both AUC and F1 score across all replicate conditions. Additional evaluations using *D*_max_ and *OR*_max_ cutoff combined with *q*-values further confirmed that the relative performance trends in the allohexaploid simulations mirrored those in the allotetraploid setting (HOBIT+ in Fig. **S6**).

**Fig. 2.**
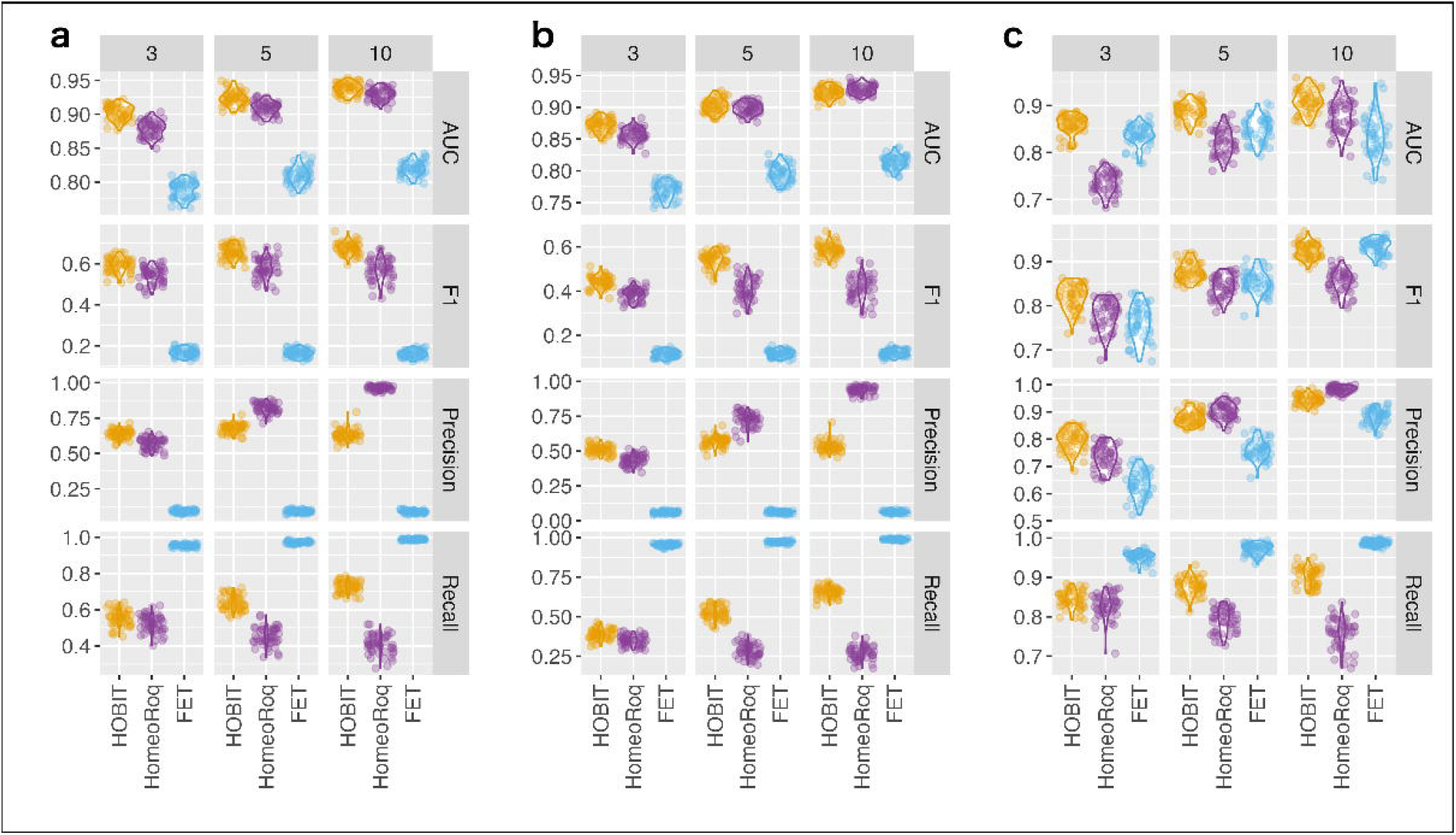
Performance metrics based on 50 artificial datasets simulating an allohexaploid species. Metrics were calculated from (**a**) all homeologs, (**b**) homeologs expressed across all subgenomes and conditions, and (**c**) homeologs with at least one subgenome unexpressed under one of the conditions.

When comparing the allotetraploid and allohexaploid simulations, HOBIT maintained consistent AUC and F1 score across all replicate conditions. In contrast, HomeoRoq showed improved performance in the allohexaploid setting, narrowing the performance gap with HOBIT in both AUC and F1 score. Conversely, FET exhibited reduced AUC and F1 score across all replicate numbers.

### Evaluation of HOBIT using alternative estimation strategies

Three HOBIT variants were evaluated using artificial datasets simulating allotetraploids and allohexaploids, following the same evaluation protocol as the default HOBIT implementation. The results from both simulations showed consistent trends. Specifically, replacing the NB distribution with the ZINB distribution showed no clear advantage (HOBITZ in Fig. **S2**–**S6**). Similarly, drawing 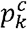 from the Dirichlet prior distribution instead of the NIP distribution showed no noticeable benefit (HOBITD in Fig. **S2**–**S6**). Furthermore, simulation studies revealed that estimating *ϕ* ^*c*^ within the MCMC sampling often led to infinite values for many homeologs, resulting in convergence failures.

### Effect of normalizing log-likelihood differences in LRT

To assess the impact of log-likelihood difference normalization during the LRT, we further evaluated the performance of HOBIT using *p*_raw_-values for AUC calculation in both allotetraploid and allohexaploid simulated datasets. The results were consistent across both dataset types (HOBITr in Fig. **S2**–**S6**). As expected, AUC calculated using *p*_raw_-values were identical to that computed using the *p*-values. This is because, although the normalization reduces the magnitude of the likelihoods during the LRT, it does not alter their relative ranking. Therefore, the relative order of *p*-values remains theoretically identical to that of the *p*_raw_-values, causing identical AUC. In contrast, when comparing *q*_raw_-values (computed from *p*_raw_-values) to the *q*-values, we found that the use of *q*_raw_-values substantially increased recall but markedly reduced precision. This trade-off became more pronounced as the number of replicates increased, resulting in a decline in F1 score. As a result, although no significant difference in F1 score was observed between *q*-values and *q*_raw_-values in datasets with three replicates, the advantage of using *q*-values became evident in datasets with five and ten replicates.

### Execution time

The average execution times for analysing a dataset simulating an allotetraploid species with three replicates were 16.0 minutes for HOBIT, 8.3 minutes for HomeoRoq, and 45 seconds for FET. Although execution time varied with the number of replicates, the impact was minimal. On average, HOBIT required 16.5 minutes for datasets with five replicates and 18.7 minutes for those with ten replicates. In comparison, HomeoRoq took 8.2 and 8.2 minutes each, and FET completed the analyses in 64 and 110 seconds, respectively.

The analysis of datasets simulating allohexaploid species required more computational time for all three methods. On average, HOBIT took 16.8, 18.5, and 25.9 minutes to process datasets with three, five, and ten replicates, respectively. FET completed the analyses more quickly, similar to its performance on allotetraploid simulations, taking 55, 82, and 129 seconds for the same replicate counts. In contrast, HomeoRoq required 25.4, 25.4, and 25.4 minutes, which is approximately three times longer than its execution times for the corresponding allotetraploid datasets. This increase in runtime for HomeoRoq is primarily due to the one-vs-rest strategy applied in the allohexaploid setting, which involves three pairwise comparisons per dataset instead of a single comparison as in the allotetraploid case.

### Ratio-shifted homeologs between habitats in C. flexuosa

When applied to the TMM-normalized count data of the *C. flexuosa* dataset, HOBIT identified 46, 124, and 130 RSHs between the two habitats on April 18, May 2, and May 16, respectively. Using FPKM-normalized data, HOBIT detected 149, 136, and 132 RSHs on the same dates. In comparison, HomeoRoq using FPKM identified substantially more RSHs, reporting 603, 436, and 353 on April 18, May 2, and May 16, respectively. Full lists of RSHs detected by each approach are available in Data S1.

HOBIT using TMM-normalized counts identified fewer RSHs on April 18 compared to later dates, while HomeoRoq detected a larger number on this date. A detailed inspection revealed that the absence of one biological replicate at the dry habitat on April 18 caused both methods to predominantly identify RSHs with extreme subgenome bias, where the HER of the A-subgenome in the dry habitat approached either 0 or 1 (Fig. **3a**). In contrast, on May 2 and May 16, both methods detected RSHs with a broader range of HERs, including more balanced expression across subgenomes.

**Fig. 3.**
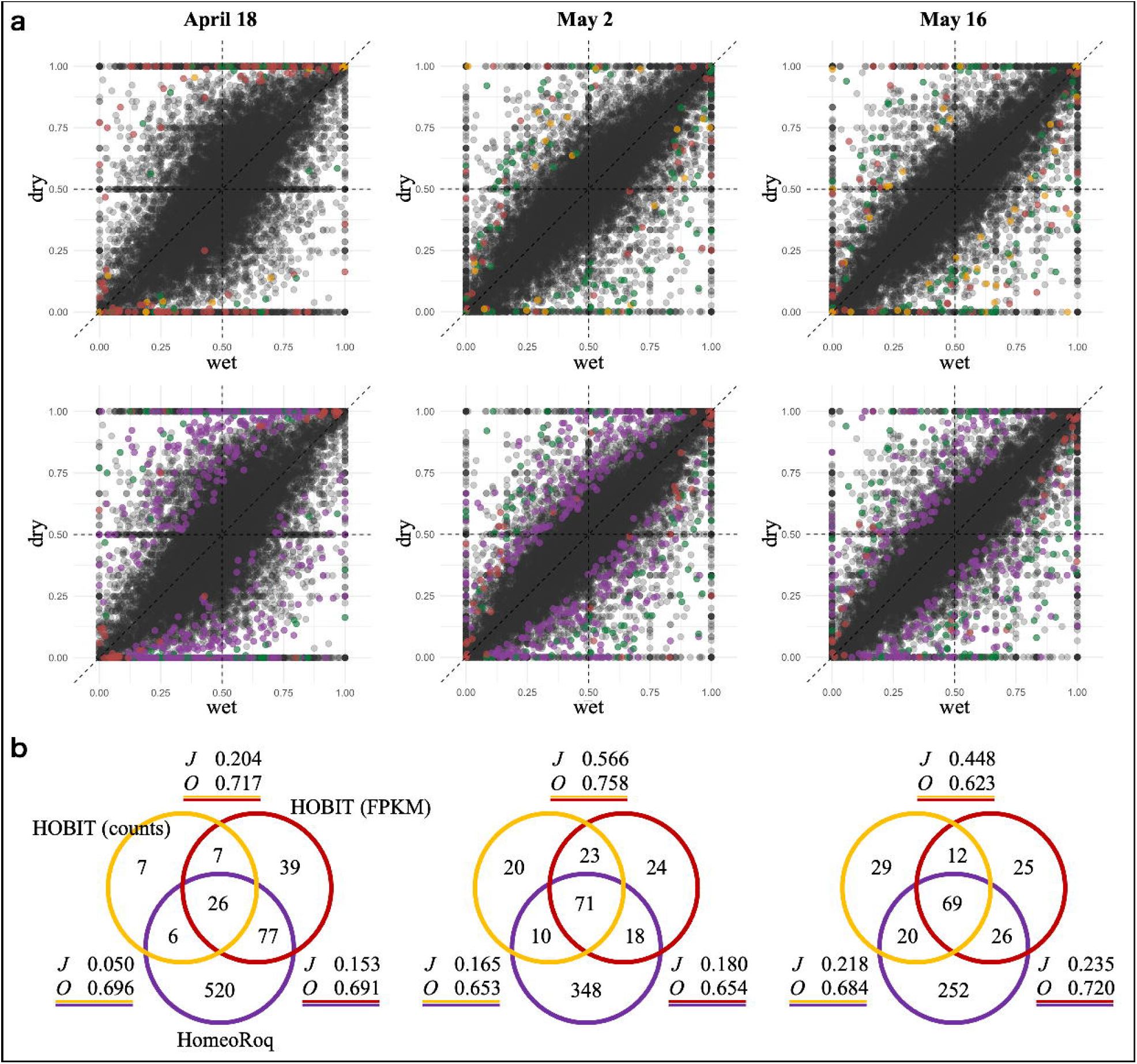
Ratio-shifted homeologs (RSHs) in *Cardamine flexuosa* detected between two distinct ecological habitats. (**a**) Scatter plots showing homeolog expression ratios for the subgenome derived from *Cardamine amara* comparing dry and wet habitats under three different sampling dates. The top row shows RSHs uniquely identified by HOBIT using count data (orange), HOBIT using FPKM (red), and those shared by both methods (green). The bottom row displays RSHs uniquely identified by HOBIT using FPKM (red), HomeoRoq (purple), and both (green). (**b**) Venn diagrams illustrating the overlap of RSHs identified by HOBIT using TMM-normalized counts (orange), HOBIT using FPKM (red), and HomeoRoq using FPKM (purple). Values after *J* and *O* represent the Jaccard index and overlap coefficients between any two RSH sets, respectively.

An overlap analysis was conducted to compare RSHs identified by the three approaches. Aside from April 18, which was affected by the missing replicate, the other two dates showed consistent overlap patterns characterized by low Jaccard indices but high overlap coefficients across all pairwise comparisons (Fig. **3b**). For instance, the comparison between HOBIT using TMM-normalized counts and HomeoRoq yielded Jaccard indices ranging from 0.169 to 0.229 and overlap coefficients between 0.653 and 0.692. The low Jaccard indices were primarily due to the substantially larger number of RSHs detected by HomeoRoq (i.e., 436 and 353) compared to HOBIT (i.e., 136 and 132). In contrast, the high overlap coefficients indicate that over 65% of RSHs identified by HOBIT were also detected by HomeoRoq.

GO enrichment analysis was performed for the RSHs identified by each approach. A large number of GO terms were enriched in the analysis using HomeoRoq, consistent with its identification of a larger number of RSHs. GO:0006865 (amino acid transport), which is known to contribute to osmotic response, as well as GO terms related to defence, were enriched across most analyses. On May 16, GO:0009414 (response to water deprivation) was enriched in the results from both HOBIT using TMM-normalized counts and HomeoRoq. GO:0009737 (response to abscisic acid) was significantly enriched only among RSHs detected by HomeoRoq, reflecting the larger number of RSHs. The completed GO enrichment analysis results are available in Data S2.

### Ratio-shifted homeologs responding to submergence stress in C. insueta

Among the 19,654 highly expressed homeolog pairs, 288 (1.47%) were identified as RSHs, homeologs that shifted their expression ratios at least at one of the nine time points following submergence, by using HOBIT (Data S3). Focusing on RSHs with orthologs in *A. thaliana*, we found that they were associated with key biological functions, including photosynthesis, stress response, energy metabolism, and homeostasis, and exhibited significant HER shifts after submergence treatment (Fig. **4**).

**Fig. 4.**
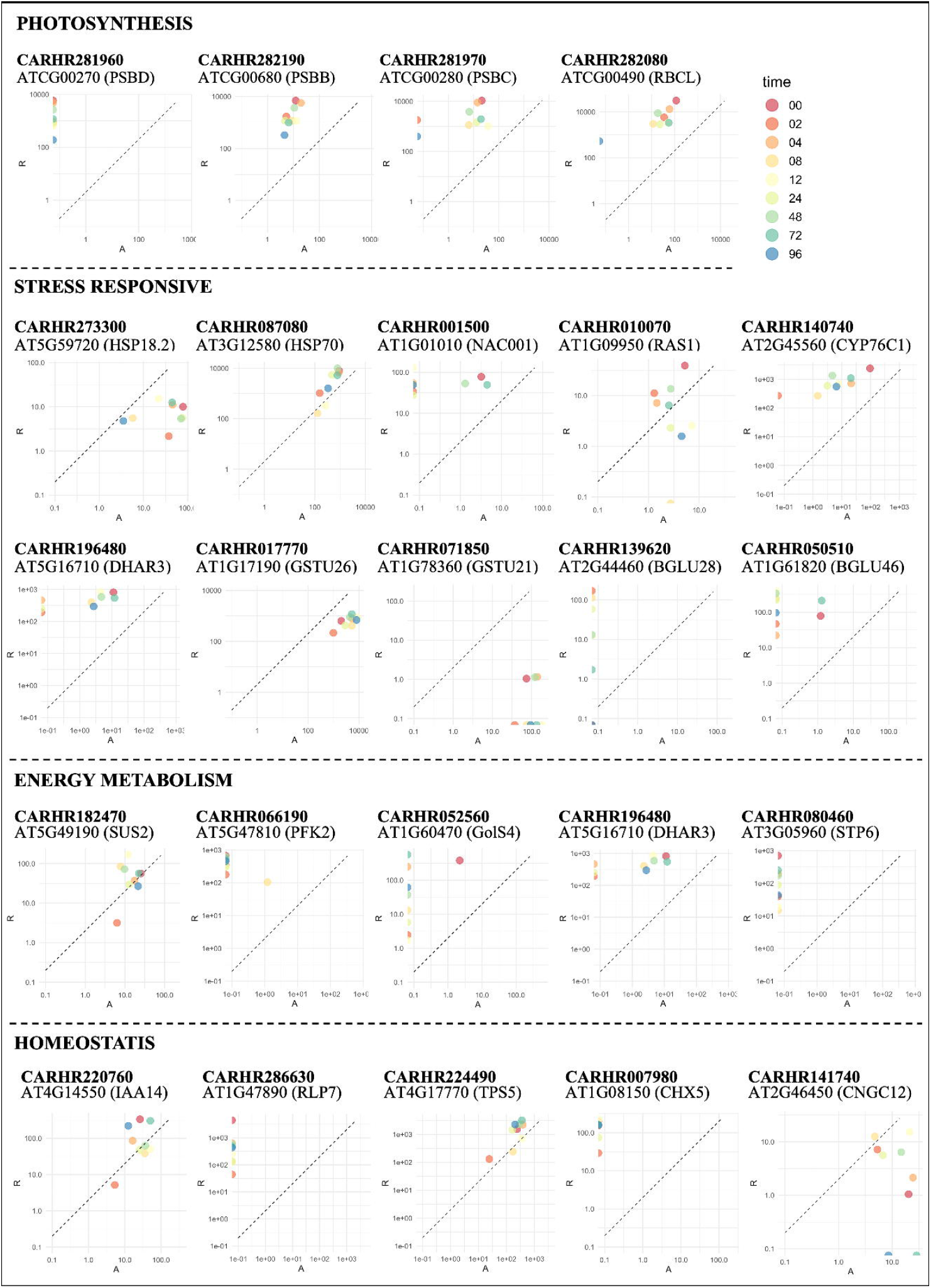
Homeolog expression dynamics in *Cardamine insueta* following submergence. Scatter plots display homeolog expression levels from the subgenome derived from *Cardamine amara* (*x*-axis) and *Cardamine rivularis* (*y*-axis) across nine time points after submergence. The dashed line (*y* = 2*x*) represents the expected expression balance based on the genomic composition, reflecting the presence of one subgenome from *C. amara* and two from *C. rivularis*.

RSHs related to photosynthesis were strongly downregulated in both subgenomes after submergence. Among them, *CARHR281960* (*PSBD*), *CARHR281970* (*PSBC*), and *CARHR282080* (*RBCL*) were suppressed in the A-subgenome at 96 h after submergence. RSHs related to stress responses were also downregulated but exhibited diverse expression patterns. Homeologs such as *CARHR087080* (*HSP70*) and *CARHR140740* (*CYP76C1*) were down-regulated in both subgenomes. Other homeologs, including *CARHR273300* (*HSP18*.*2*) and *CARHR017770* (*GSTU26*), were up- or down-regulated only from the A-subgenome, while maintaining stable expression in the R-subgenome. Additionally, homeologs such as *CARHR010070* (*RAS1*), *CARHR196480* (*DHAR3*), and *CARHR139620* (*BGLU28*) were predominantly expressed in the R-subgenome before submergence and showed reduced expression after treatment.

Expanding the analysis beyond the functionally annotated homeologs revealed additional diverse HER patterns (Fig. **S7**). A prominent pattern involved homeologs primarily expressed from one subgenome, with the HER shifting after submergence. For example, *CARHR103390* (*AT2G05810*) and *CARHR112990* were predominantly expressed in the R-subgenome and showed decreased expression after submergence. In contrast, *CARHR009470* (*AT1G09500*) and *CARHR070320* (*IGMT5*) were also R-subgenome-dominant but showed increased expression. Similarly, *CARHR057200* and *CARHR138680*, which were mainly expressed from the A-subgenome, exhibited reduced expression after submergence. Another pattern involved homeologs expressed in both subgenomes, where expression remained stable in one subgenome but changed in the other, leading to HER shifts over time. Examples of homeologs that changed their expression in the A-subgenome, while remaining stable in the R-subgenome, include *CARHR091610* (*NSP1*), *CARHR093140* (*PAP17*), *CARHR281870* (*ATPA*), and *CARHR147260* (*CYP71B24*). In comparison, homeologs that changed in the R-subgenome while maintaining stable expression in the A-subgenome included *CARHR095340* (*NTMC2T5*.*2*), *CARHR071710* (*AT1G78260*), and *CARHR189390*.

GO enrichment analysis confirmed that the RSHs were significantly associated with submergence stress responses, including GO:0015979 (photosynthesis), GO:0019684 (photosynthesis and light reaction), GO:0042542 (response to hydrogen peroxide), GO:0005982 (starch metabolic process), GO:0005986 (sucrose biosynthetic process), and GO:0042631 (cellular response to water deprivation). These enriched GO terms reflected the stress response to the submergence treatment, in agreement with their experimental condition.

### Ratio-shifted homeologs between tissues and developmental stages in wheat

A total of 48 (0.36%) and 105 (0.77%) homeolog triads were identified as RSHs between developmental stages (W2.0 vs. W3.5) in the leaf and shoot apex, respectively (Data S4). In the leaf, RSHs were enriched for homeologs associated with stress responses. For example, *TraesCS4A02G092900* (heat shock protein), *TraesCS6A02G047600* (peroxidase), *TraesCS2A02G573400* (peroxidase), and *TraesCS7A02G191200* (aquaporin) exhibited shifts in HERs, reflecting differential homeolog utilization under changing physiological conditions (Fig. **5a**).

**Fig. 5.**
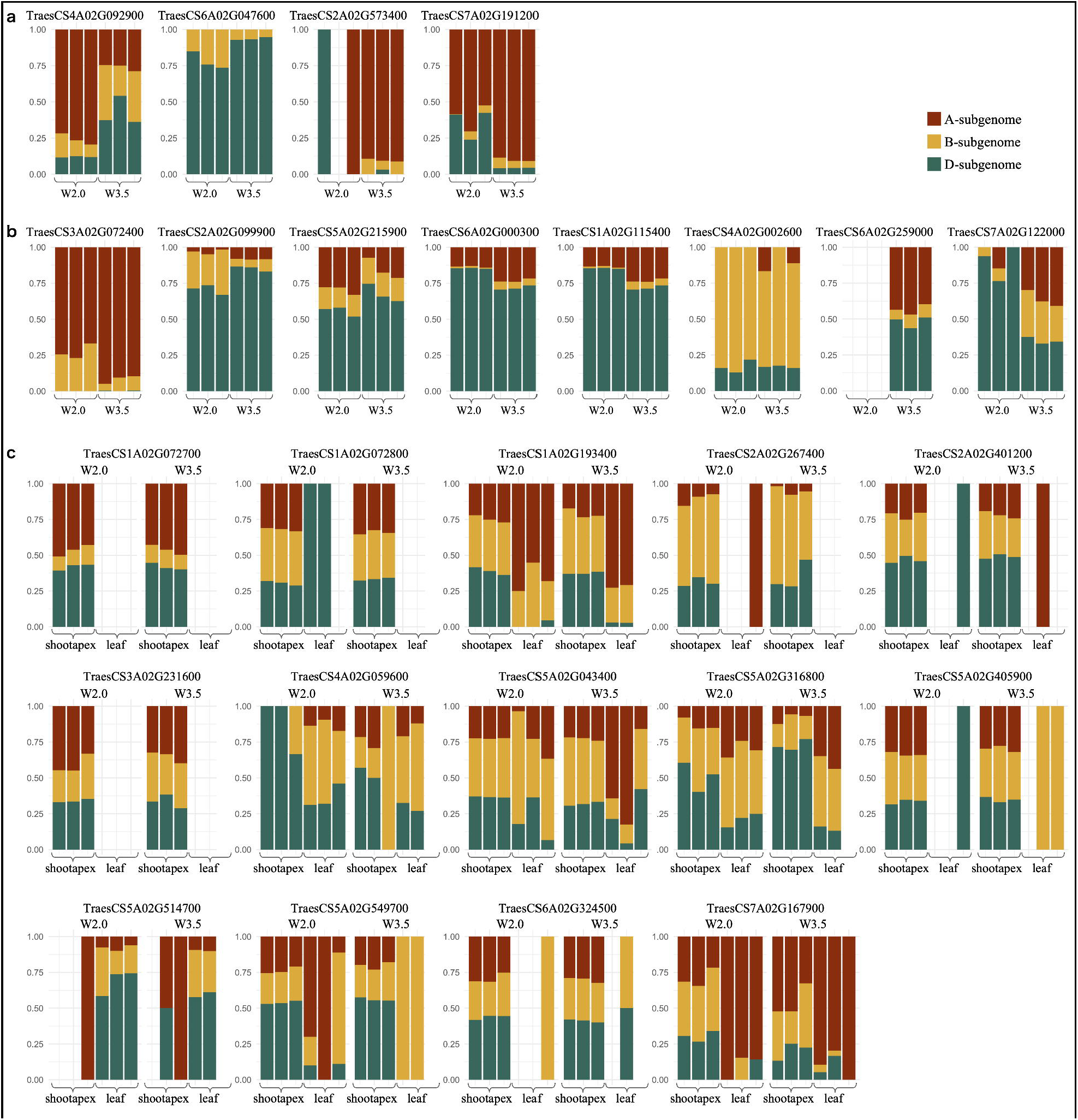
Visualization of homeolog expression ratios (HERs) for ratio-shifted homeologs (RSHs). Bar charts display HERs of the three subgenomes under different conditions. Absence of a bar indicates that the homeolog was not expressed from all three subgenomes. The colour composition of each bar reflects the HER computed from normalized RNA-Seq data. (**a**) Examples of RSHs detected between W2.0 and W3.5 in leaf tissue. (**b**) Examples of RSHs detected between W2.0 and W3.5 in shoot apex. (**c**) Examples of homeobox genes detected as RSHs between leaf and shoot apex across both developmental stages.

In the shoot apex, homeologs related to developmental processes were frequently observed among the RSHs. Several homeologs associated with ethylene signalling, a key plant hormone regulating plant growth and development, showed dynamic HER shifts (Fig. **5b**). For instance, *TraesCS3A02G072400* (E3 ubiquitin-protein ligase) was predominantly expressed from the A-subgenome, followed by the B-subgenome, with scarce expression from the D-subgenome. The HER of the A-subgenome increased at the W3.5 stage compared to W2.0. In contrast, *TraesCS2A02G099900* (ethylene insensitive 3) and *TraesCS5A02G215900* (ethylene-responsive transcription factor) were dominantly expressed from the D-subgenome, with small contribution from the A- and B-subgenomes; both exhibited increased HERs of the D-subgenome at W3.5 stage. Homeologs directly associated with development regulation also showed biased expression. *TraesCS6A02G000300* (SKP1-like protein) and *TraesCS1A02G115400* (embryo-specific 3) were dominantly expressed from the D-subgenome, followed by moderate expression from the A-subgenome and minimal expression from the B-subgenome. At stage W3.5, the HER of the A-subgenome increased, while that of the D-subgenome decreased. MADS-box transcription factor homeologs, including *TraesCS4A02G002600, TraesCS6A02G259000*, and *TraesCS7A02G122000*, also demonstrated complex HER shift patterns.

When focusing on RSHs between tissues (leaf vs. shoot apex), a substantial number of RSHs were identified: 4,694 (32.45%) and 2,662 (18.30%) homeolog triads at W2.0 and W3.5, respectively (Data S5). Among these, 1,893 RSHs were shared between the two developmental stages, indicating permanent HER differences between tissues. A manual inspection of these RSHs revealed that a subset was functionally associated with plant hormone signalling, stress responses, and developmental regulation. Specifically, RSHs included four triads related to abscisic acid, nine to auxin, four to cytokinin, ten to ethylene, and five to gibberellin. Additionally, 33 RSHs were related to F-box proteins and 13 to peroxidases, both of which are implicated in stress responses. Moreover, 10 RSHs were associated with embryogenesis or the flowering process, 14 with homeobox genes, and three with MADS-box transcription factors. Focusing on homeobox homeologs, the majority were preferentially expressed in the shoot apex and suppressed in the leaf across both developmental stages, with several exception such as *TraesCS5A02G514700* (Fig. **5c**).

## Discussion

### Inputs of HOBIT

HOBIT performs an LRT for each homoeologous set independently, assuming that the count data follow a NB distribution. The NB distribution was chosen for its ability to handle overdispersion, which is a common characteristic of RNA-Seq data (Robinson & Smyth, 2007; Love *et al*., 2014; Chen *et al*., 2025). Consequently, HOBIT accepts any type of normalized count data that retains the fundamental properties of the count data and enables comparisons across subgenomes and experimental conditions.

However, it is important to emphasize that the choice of normalization method can affect RSH detection. As demonstrated in the *C. flexuosa* case study, HOBIT produced differing sets of RSHs when applied to TMM-normalized counts versus FPKM (Fig. **3**). Jaccard indexes indicate that approximately 50% of the RSHs identified using one normalization method were not detected using the other, underscoring the influence of normalization on RSH detection.

Because the optimal normalization method can vary depending on factors such as the experimental design, read mapping strategy, reference genome quality, and analytical objectives, HOBIT does not recommend a single best normalization approach. While TMM and FPKM have been applied in case studies, other methods such as mean-of-ratio normalization (Anders & Huber, 2010) and transcripts per million (TPM) (Li *et al*., 2010) are also compatible with HOBIT. However, caution is advised when using normalization methods that adjust the total library size, such as FPKM and TPM, because they may distort expression distributions when transcript abundance distributions differ across samples, potentially leading to spurious results (Zhao *et al*., 2020, 2021). Therefore, researchers are encouraged to carefully select a normalization method that suits the characteristics of their datasets.

### Outputs of HOBIT

HOBIT outputs statistical metrics for each homeolog, including: (i) *p*_raw_-value, derived directly from the LRT; (ii) *p*-value, derived from the LRT with log-likelihoods normalized by the number of replicates; (iii) *q*-value, which adjusts the *p*-value for multiple testing; (iv) *D*_max_, the maximum absolute HER shifts among conditions observed in at least one subgenome, computed from posterior samples; and (v) *OR*_max_, the maximum odds ratio of HER shifts, also computed from posterior distributions.

The *p*_raw_-value serves as a standard measure of statistical significance in LRT. However, it is highly sensitive to the number of replicates (Wilks, 1938; Dziak *et al*., 2014; Gudicha *et al*., 2017). This sensitivity arises because the likelihood increases with the number of observations, which can amplify even minor differences between the competing models. As a result, LRT conducted with larger number of replicates (e.g., five or ten) can yield statistical significance that does not reflect biologically meaningful differences. To address this issue and reduce false positives caused by sample size effects, we applied an ad hoc normalization that adjusts the log-likelihood difference according to the number of replicates. Simulation-based evaluations demonstrated that F1 scores were the highest when detecting RSHs based on *q*-values calculated from the normalized *p*-values, consistently outperforming the results based on *p*_raw_-values across varying replicate conditions (HOBIT and HOBITr in Fig. **S2**–**S6**). These findings highlight that the normalization procedure offers a more robust approach for detecting HER shifts, particularly in datasets with larger sample sizes.

Moreover, to enhance interpretability in a biological context, incorporating *D*_max_ and *OR*_max_ alongside *q*-values can help prioritize RSHs with not only statistical but also practical relevance (HOBIT+ in Fig. **S2**–**S6**). A commonly recommended threshold might include *q*-value <0.01 in combination with *D*_max_ >0.2 or *OR*_max_ >2.0, akin to practices in differential expression analysis where a fold-change cutoff complements *q*-value thresholds. This integrated approach balances statistical stringency with biological meaning, making HOBIT a robust and practical framework for analysing HERs in real-world allopolyploid datasets.

### Parameters of HOBIT

RNA-Seq count data in some cases contain a large proportion of zero counts. To account for this, some studies have modelled such data using zero-inflated distributions, such as zero-inflated Poisson and ZINB distributions, which distinguish between structural zeros (genes not expressed) and sampling zeros (zeros due to random variation). Given that certain homeologs may be completely suppressed under specific conditions, causing a large proportion of zero counts, we also evaluated HOBIT implementation using ZINB distribution. However, the results showed no clear advantage of ZINB over NB distributions (HOBITZ in Fig. **S2**–**S6**). One possible reason may be related to the actual characteristics of RNA-Seq read counts, as some studies have shown that the NB distribution is sufficiently robust for modelling RNA-Seq (Svensson, 2019; Zhao *et al*., 2022).

Regarding dispersion (*ϕ*^*c*^) estimation, it is known that dispersions share information across all homeologs (Robinson & Smyth, 2007; Love *et al*., 2014), making independent estimations for each homeolog less accurate. To address this, HOBIT first estimates *ϕ*^*c*^ across all homeologs using edgeR (Chen *et al*., 2025) before performing the MCMC sampling. Simulation studies confirmed that pre-computing dispersions stabilized MCMC convergence compared to estimating dispersions independently for each homeolog via MCMC sampling.

### Biological findings of HOBIT

Analysis of real RNA-Seq data from multiple types of allopolyploids, which have different degrees of dispersions, demonstrated the effectiveness of HOBIT in detecting biologically meaningful shifts in HERs. In a case study of *C. flexuosa*, the majority of RSHs detected by HOBIT were also identified by HomeoRoq (Fig. **3**), indicating that both algorithms can capture a consistent set of RSHs. Similar categories were identified by GO enrichment analysis, although the statistical significance in HOBIT was limited due to the small number of RSHs. Further application to *C. insueta* successfully revealed dynamic HER changes across a time series following the submergence treatment (Fig. **4** and Fig. **S7**), highlighting the robustness and adaptability of HOBIT in allopolyploids with an uneven number of subgenomes across multiple conditions. Moreover, when applied to an allohexaploid species, wheat, HOBIT uncovered a notable trend; HERs differed more substantially between tissues than between developmental stages. Thus, many homeolog triads exhibit tissue-specific HERs, whereas relatively few exhibit HER shifts across developmental stages. This suggests that while different tissues preferentially utilize different homeologs, HERs tend to remain stable once established, regardless of developmental stage. These findings demonstrate the utility of HOBIT in extracting novel biological insights even from published datasets. Although further experimental validation and broader analyses are required, HOBIT offers a promising framework for exploring patterns of homeolog regulation and environmental adaptation in the allohexaploid, wheat.

### Advantages and disadvantages of HOBIT

Assuming that count data follow a NB distribution, HOBIT effectively accounts for overdispersion, allowing for more flexible and accurate detection of RSHs, including cases that HomeoRoq tends to overlook. In addition, HOBIT supports RSH detection across more than two conditions, similar to ANOVA, and does not impose constraints on subgenome composition. These features make HOBIT a versatile and powerful tool for understanding how allopolyploids achieve environmental adaptability, a key aspect in the development of climate-resilient crops in the face of global environmental change.

The primary limitation of HOBIT is its longer runtime: analysing 10,000 homeolog pairs requires approximately half an hour, which is considerably slower than differential expression analysis using R packages, such as edgeR (Chen *et al*., 2025) and DESeq2 (Love *et al*., 2014), typically completed in seconds. However, because such analyses are usually performed only once per dataset in real-world applications, we suggest that an extended runtime is generally not a major concern.

Another important consideration is the consistency of results of HOBIT with those from other methods, such as HomeoRoq. A case study using the *C. flexuosa* dataset showed that HOBIT and HomeoRoq identified overlapping but different sets of RSHs, even though both methods used the same FPKM input. This discrepancy likely derives from fundamental differences in modeling conceptions. First, HOBIT assumes that homeolog expression follows a NB distribution, while HomeoRoq models counts using a hierarchical normal distribution with an additional noise term to account for overdispersion. Second, HOBIT conducts an LRT by comparing one model that assumes that HERs differ across conditions with another model that assumes that HERs are shared, while HomeoRoq estimates the probability that HERs remain unshifted between conditions based on observed homeolog expression. Given that different analysis methods often yield divergent results in differential expression analysis of RNA-Seq data (Rajkumar *et al*., 2015; Tarazona *et al*., 2015; Zambelli *et al*., 2018; Li *et al*., 2022), such discrepancies are, to some extent, unavoidable and reflect differences in methodological assumptions and modeling strategies. Therefore, when interpreting detection results, whether RSH detection or differential expression analysis in general, it is important to recognize that each method may be more sensitive to certain types of gene or homeolog changes while potentially overlooking others due to its specific methodological assumptions and modeling strategies.

## Acknowledgements

This work was supported by the Japan Society for the Promotion of Science [JP25K09719 to JSun, JP23K23582 to KKS, and JP22H05179 to JSun and KKS], the Japan Science and Technology Agency [JPMJCR16O3 to KKS and JSese], and the Swiss National Science Foundation [31003A_212551 to KKS]. We thank Wei Cao (RCAIT, NARO) for contributing to the discussion.

## Author contributions

JSun: conceptualization, supervision, investigation, data curation, methodology, formal analysis, validation, software, funding acquisition, writing – original draft, writing – review and editing. JSese: funding acquisition, writing – review and editing. KKS: formal analysis, funding acquisition, writing – original draft, writing – review and editing.

## Data availability

RNA-Seq datasets used in the case studies were downloaded from the NCBI SRA database under the following accession numbers: DRA006316 for *Cardamine flexuosa*, DRA009830 for *Cardamine insueta*, and SRA1141904 for wheat. Homeolog expression data quantified from these RNA-Seq datasets, along with analysis scripts supporting the findings of this study, are available on Zenodo under the identifier 10.5281/zenodo.15259060.

## Supplementary data

SI_Methods: A PDF file containing supplementary methods (Methods S1–S3).

SI_Figures: A PDF file containing supplementary figures (Figures S1–S7).

SI_Data: A ZIP archive containing supplementary data in Excel format (Data S1–S5).

