## Supplementary material for "A likelihood ratio test for detecting shifts in homeolog expression ratios in allopolyploids": SI Figures

### Figure S1

**Overview of the process for generating artificial RNA-Seq read counts in allopolyploids.** (**A**) A population of mean and variance values is derived from real RNA-Seq read counts, and a model fitting log-transformed variance as a function of log-transformed mean is constructed. (**B**) A specified number of mean values are sampled from this population. Random values drawn from a 1 + Gamma distribution are applied to these sampled means to generate means for the control and treatment conditions, introducing a small proportion of differentially expressed genes. (**C**) The proportion of relative homeolog expression and small expression changes between conditions are generated using random values sampled from normal and Gamma distributions, introducing biasedly expressed homeologs. (**D**) A small proportion of means are randomly set to zero to mimic unexpressed homeologs. (**E**) The adjusted means are used to sample count data from a negative binomial distribution, using the sampled means and the fitted variance values.

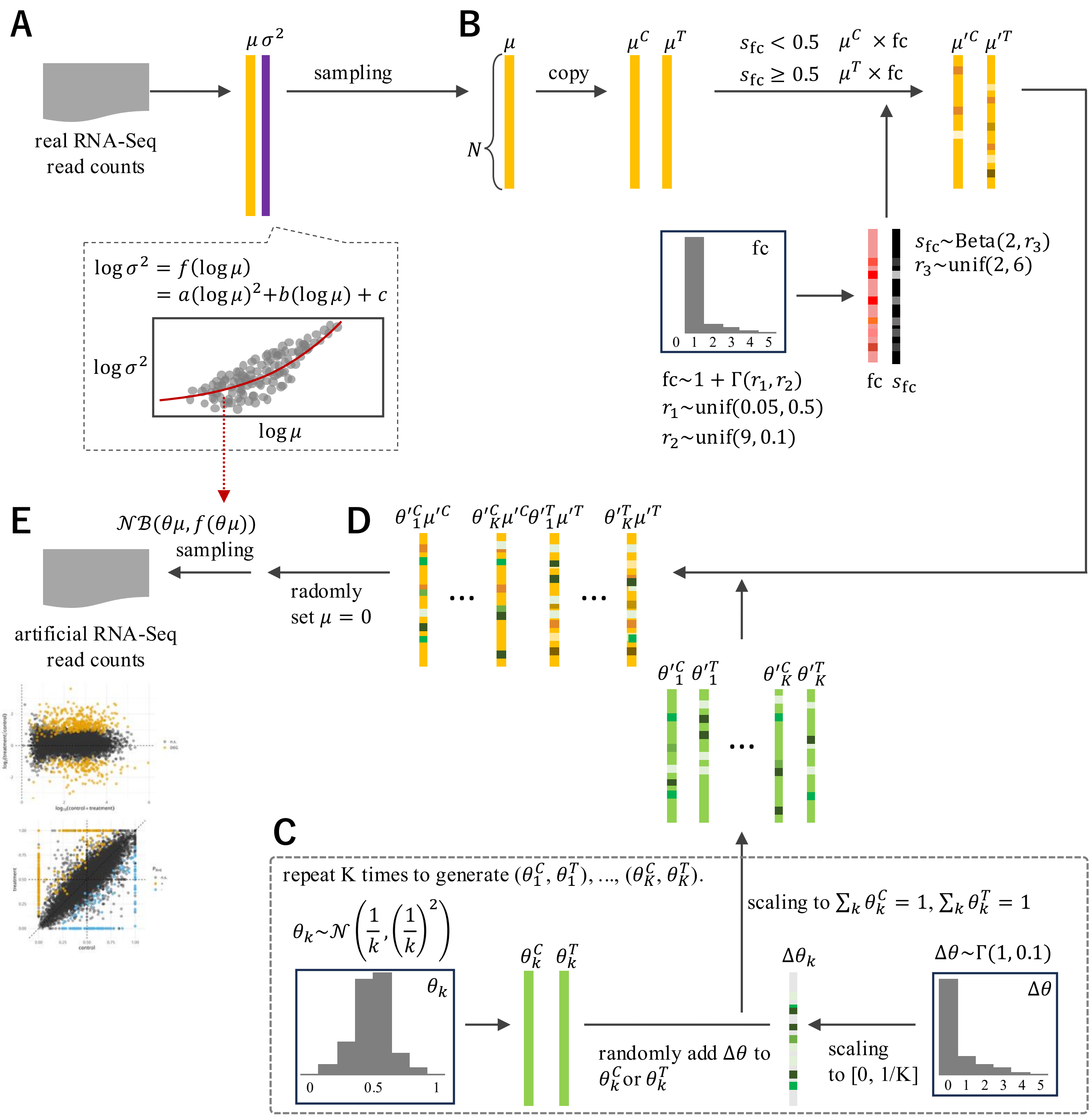

### Figure S2

**Performance metrics on artificial datasets simulating an allotetraploid.** Violin plots with overlaid scatter points show the performance metrics (AUC, F1 score, precision, and recall) across datasets with different number of replicates (3,5, and 10) for HOBIT, its variations, HomeoRoq, and Fisher's exact test (FET). HOBIT indicates the default configuration, while HOBITD uses a Dirichlet prior for modeling homeolog expression ratios and HOBITZ employs a zero-inflated negative binomial (ZINB) distribution for modeling expression levels. Method names with the suffix "r" indicate that performance metrics were calculated using $p_{raw}$-values, while those with the suffix "+" reflect results after applying additional cutoff thresholds ($D_{max}$ >0.2 or $OR_{max}$ >2.0) to the $q$-values (or $q_{raw}$-values for "r+"). Metrics were computed under three scenarios, (**A**) using all homeologs, (**B**) using only homeologs expressed across all subgenomes and conditions, and (**C**) using homeologs with no expression in at least one subgenome under one condition. Ground-truth ratio-shifted homeologs in artificial datasets were defined as $D_{max}$ >0.2 or $OR_{max}$ >2.0, where $D_{max}$ and $OR_{max}$ were obtained from simulation condition.

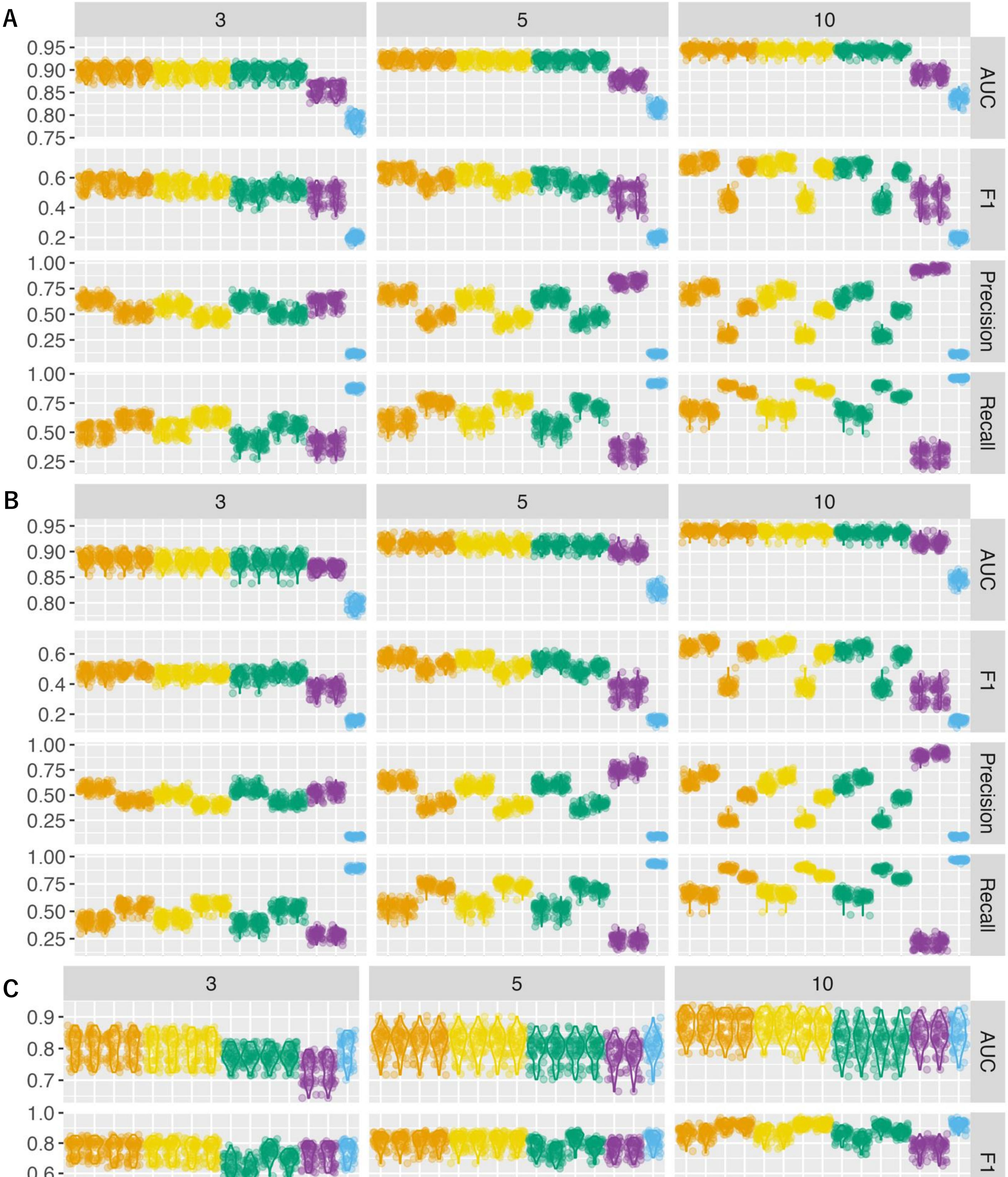

### Figure S3

**Performance metrics on artificial datasets simulating an allotetraploid.** Violin plots with overlaid scatter points show the performance metrics (AUC, F1 score, precision, and recall) across datasets with different number of replicates (3,5, and 10) for HOBIT, its variations, HomeoRoq, and Fisher's exact test (FET). HOBIT indicates the default configuration, while HOBITD uses a Dirichlet prior for modeling homeolog expression ratios and HOBITZ employs a zero-inflated negative binomial (ZINB) distribution for modeling expression levels. Method names with the suffix "r" indicate that performance metrics were calculated using $p_{\mathrm{raw}}$-values, while those with the suffix "+" reflect results after applying additional cutoff thresholds ($D_{\max}$ >0.2 or $OR_{\max}$ >2.0) to the $q$-values (or $q_{\mathrm{raw}}$-values for "r+"). Metrics were computed from all homeologs. Ground-truth ratio-shifted homeologs in artificial datasets were defined as $D_{\max}$ >0.2 or $OR_{\max}$ >1.5, where $D_{\max}$ and $OR_{\max}$ were obtained from simulation condition.

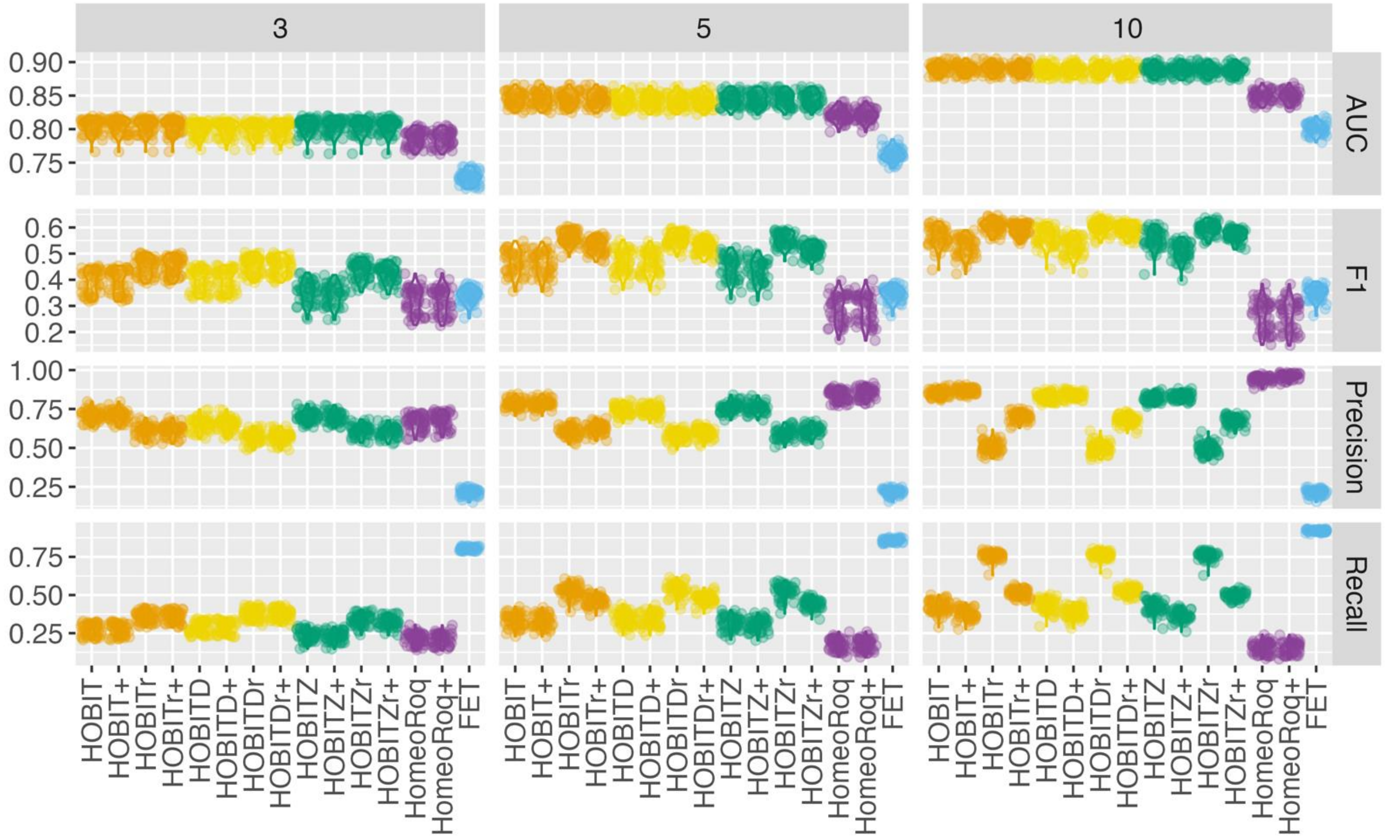

### Figure S4

**Performance metrics on artificial datasets simulating an allotetraploid.** Violin plots with overlaid scatter points show the performance metrics (AUC, F1 score, precision, and recall) across datasets with different number of replicates (3,5, and 10) for HOBIT, its variations, HomeoRoq, and Fisher's exact test (FET). HOBIT indicates the default configuration, while HOBITD uses a Dirichlet prior for modeling homeolog expression ratios and HOBITZ employs a zero-inflated negative binomial (ZINB) distribution for modeling expression levels. Method names with the suffix "r" indicate that performance metrics were calculated using $p_{\mathrm{raw}}$-values, while those with the suffix "+" reflect results after applying additional cutoff thresholds ($D_{\max}$ >0.2 or $OR_{\max}$ >2.0) to the $q$-values (or $q_{\mathrm{raw}}$-values for "r+"). Metrics were computed from all homeologs. Ground-truth ratio-shifted homeologs in artificial datasets were defined as $D_{\max}$ >0.2 or $OR_{\max}$ >3.0, where $D_{\max}$ and $OR_{\max}$ were obtained from simulation condition.

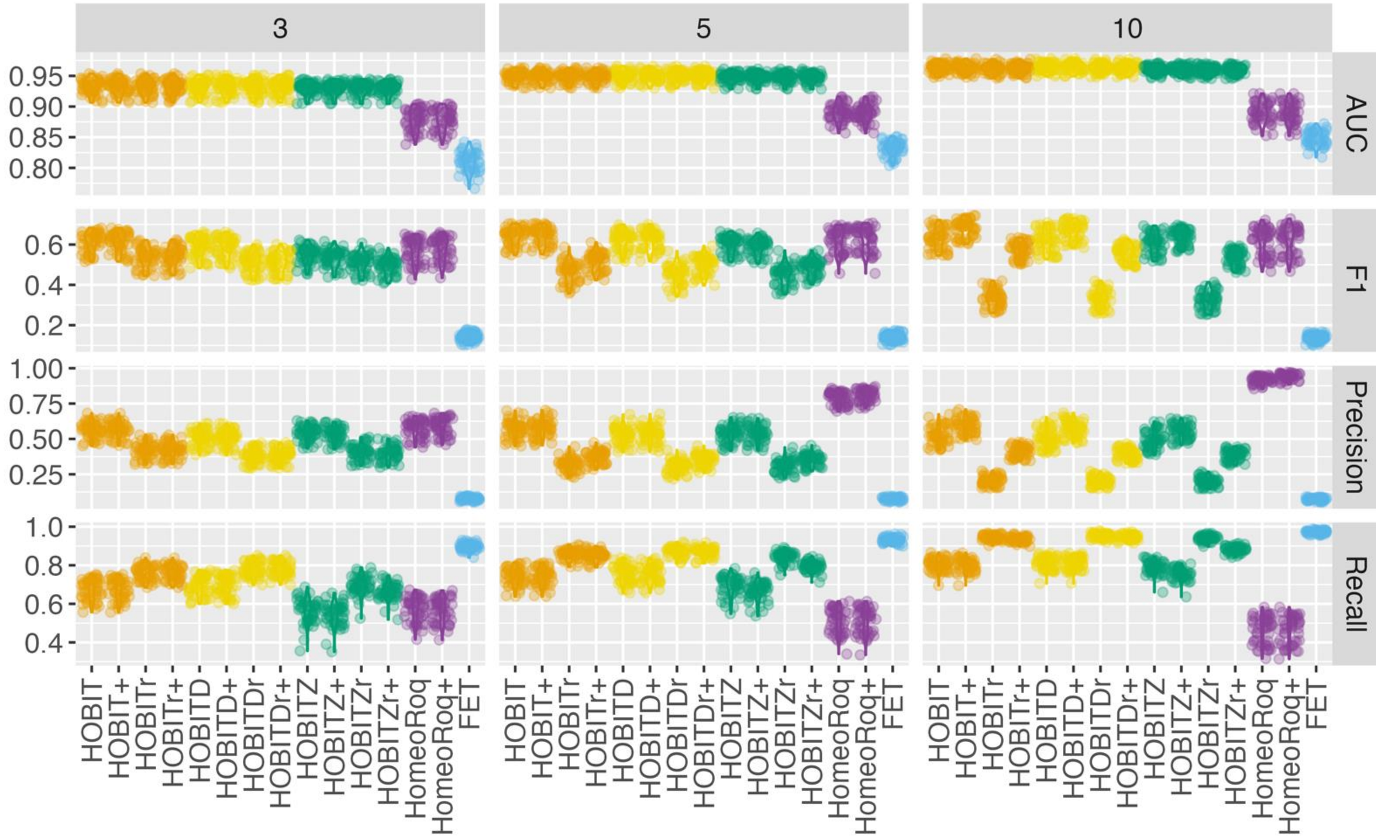

### Figure S5

**Performance metrics on artificial datasets simulating an allotetraploid.** Violin plots with overlaid scatter points show the performance metrics (AUC, F1 score, precision, and recall) across datasets with different number of replicates (3,5, and 10) for HOBIT, its variations, HomeoRoq, and Fisher's exact test (FET). HOBIT indicates the default configuration, while HOBITD uses a Dirichlet prior for modeling homeolog expression ratios and HOBITZ employs a zero-inflated negative binomial (ZINB) distribution for modeling expression levels. Method names with the suffix "r" indicate that performance metrics were calculated using $p_{\mathrm{raw}}$-values, while those with the suffix "+" reflect results after applying additional cutoff thresholds ($D_{\mathrm{max}}$ >0.2 or $OR_{\mathrm{max}}$ >3.0) to the $q$-values (or $q_{\mathrm{raw}}$-values for "r+"). Metrics were computed from all homeologs. Ground-truth ratio-shifted homeologs in artificial datasets were defined as $D_{\mathrm{max}}$ >0.2 or $OR_{\mathrm{max}}$ >3.0, where $D_{\mathrm{max}}$ and $OR_{\mathrm{max}}$ were obtained from simulation condition.

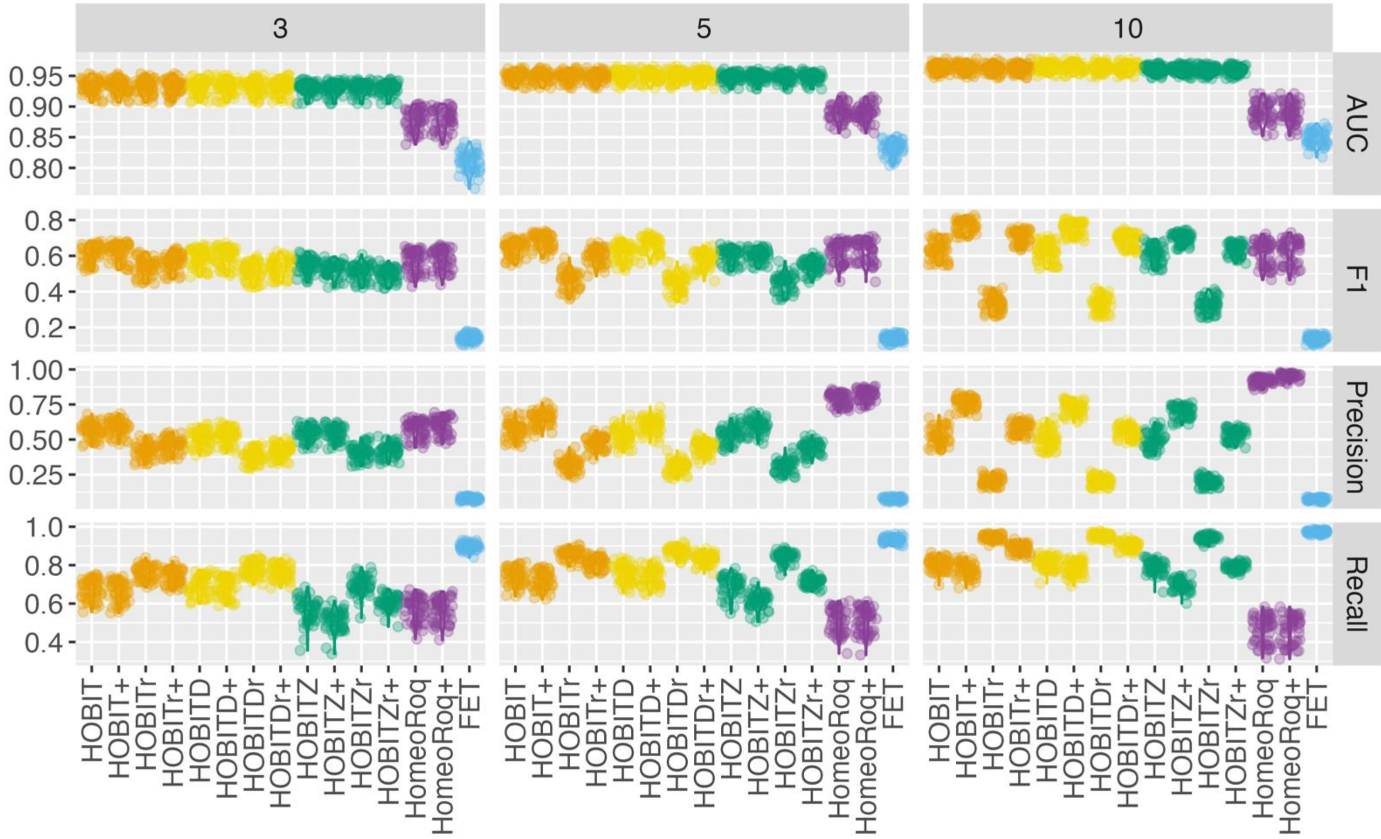

#### Figure S6

**Performance metrics on artificial datasets simulating an allohexaploid.** Violin plots with overlaid scatter points show the performance metrics (AUC, F1 score, precision, and recall) across datasets with different number of replicates (3,5, and 10) for HOBIT, its variations, HomeoRoq, and Fisher's exact test (FET). HOBIT indicates the default configuration, while HOBITD uses a Dirichlet prior for modeling homeolog expression ratios and HOBITZ employs a zero-inflated negative binomial (ZINB) distribution for modeling expression levels. Method names with the suffix "r" indicate that performance metrics were calculated using $p_{raw}$-values, while those with the suffix "+" reflect results after applying additional cutoff thresholds ($D_{max}$ >0.2 or $OR_{max}$ >2.0) to the $q$-values (or $q_{raw}$-values for "r+"). Metrics were computed under three scenarios, (**A**) using all homeologs, (**B**) using only homeologs expressed across all subgenomes and conditions, and (**C**) using homeologs with no expression in at least one subgenome under one condition. Ground-truth ratio-shifted homeologs in artificial datasets were defined as $D_{max}$ >0.2 or $OR_{max}$ >2.0, where $D_{max}$ and $OR_{max}$ were obtained from simulation condition.

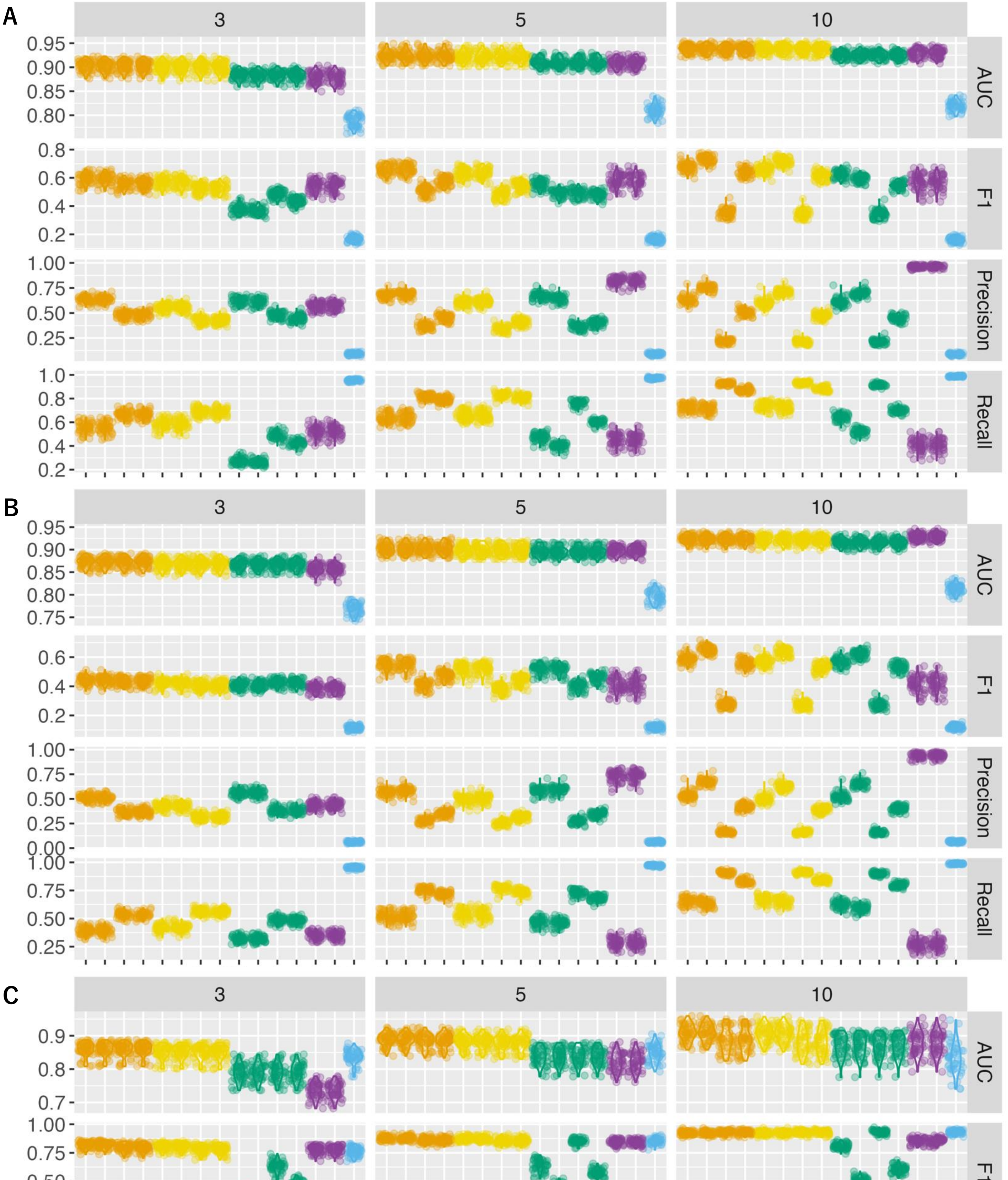

### Figure S7

**Homeolog expression dynamics in *Cardamine insueta* following submergence**. Scatter plots show homeolog expression levels from the A-subgenome (x-axis) and the R-subgenome (y-axis) across nine time points after submergence. The dashed line, $y = 2x$, represents the expected expression balance based on genomic composition, reflecting the presence of two subgenomes from *C. rivularis* and one from *C. amara*.

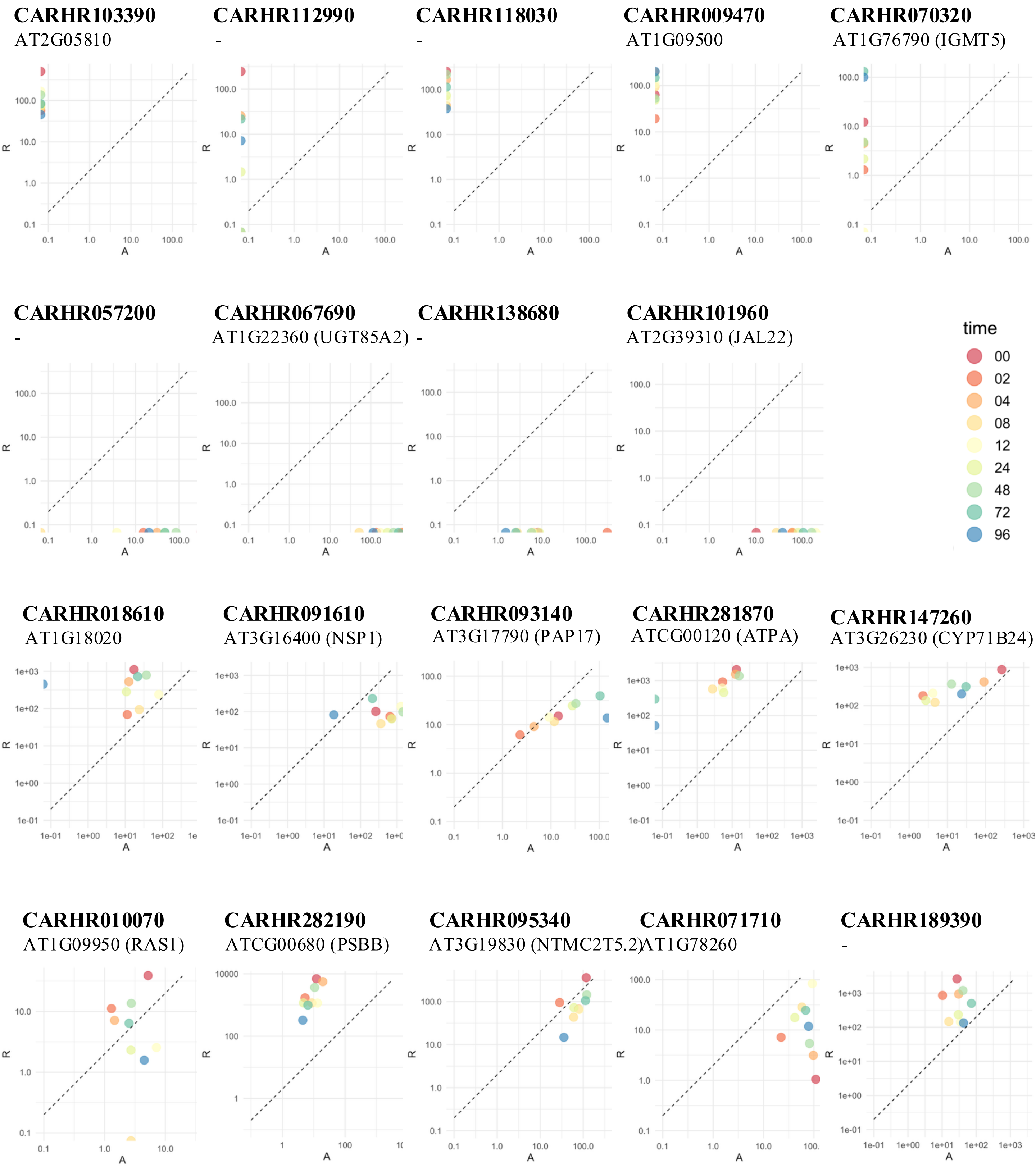
