## Supplementary material for "A likelihood ratio test for detecting shifts in homeolog expression ratios in allopolyploids": SI Methods

### Method S1: Implementing HOBIT with a zero-inflated negative binomial distribution

A zero-inflated negative binomial (ZINB) distribution has the potential to accommodate overdispersion while distinguishing between structural zeros (e.g., homeologs not expressed) and sampling zeros (e.g., zeros occurring due to random variation). The probability mass function of the ZINB distribution is given by:

$$f_{ZINB}(x|\pi,\mu,\phi) = \begin{cases} (1-\pi)f_{NB}(x|\mu,\phi), & x > 0 \\ \pi + (1-\pi)f_{NB}(x=0|\mu,\phi), & x = 0 \end{cases}$$

where $\pi$ represents the proportion of structural zeros, and $f_{NB}$ is the probability mass function of a negative binomial (NB) distribution, defined as:

$$f_{NB}(x|\mu,\phi) = \binom{x+\phi-1}{x}\left(\frac{\mu}{\mu+\phi}\right)^{x}\left(\frac{\phi}{\mu+\phi}\right)^{\phi}$$

with $\mu$ and $\phi$ represent the mean and dispersion parameters, respectively.

The test algorithm implementing ZINB follows the same steps as the NB-based approach, except for (i) the additional parameter $\pi$ is sampled from a uniform prior distribution and (ii) likelihoods are calculated using $f_{ZINB}$ instead of $f_{NB}$.

### Method S2: Sampling homeolog expression ratios from a Dirichlet prior distribution

In addition to sampling $\boldsymbol{\theta}^c$ from a uniform prior distribution with the restriction $\sum_k \theta_k^c = 1$, we also evaluated the case of sampling $\boldsymbol{\theta}^c$ is sampled from a Dirichlet prior distribution:

$$\boldsymbol{\theta}^c \sim Dirichlet(\boldsymbol{\alpha}^c)$$

where each element of $\boldsymbol{\alpha}^c$ is calculated as:

$$\alpha_k^c = \frac{K\sum_i x_{ik}^c}{\sum_k \sum_i x_{ik}^c}$$

### Method S3: Algorithm to generate artificial data

Algorithm to generate artificial RNA-Seq read count data can be roughly involved the four steps, sampling population preparation, differentially expressed homeologs (DEHs) introduction, ratio-shifted homeologs (RSHs) introduction, and counts sampling (**Figure S1**). In details, to generate one set of artificial RNA-Seq counts consisting of $N$ homeolog tuples in an allopolyploid species with $K$ subgenomes under two conditions—control (C) and treatment (T), the following steps were performed.

The mean ($\mu$) and variance ($\sigma^2$) of gene expression are first calculated from real RNA-Seq datasets to serve as a reference population (**Figure S1A**). Next, a quadratic function was fitted to describe the relationship between log-transformed mean and variance, expressed as $\log\sigma^2 = f(\log\mu)$. Then, from this reference population, $N$ of means $(\mu_1, \mu_2, \cdots, \mu_N)$ are sampled to represent the baseline expression levels of homeologs, ensuring that baseline expression is equivalent across conditions. This is formulated as $\boldsymbol{\mu}^C = \boldsymbol{\mu}^T = (\mu_1, \mu_2, \cdots, \mu_N)$, where $\boldsymbol{\mu}^C$ represents the mean expression levels of $N$ homeologs under control condition, while $\boldsymbol{\mu}^T$ represents those under the treatment condition.

Next, to introduce differentially expressed genes (DEGs) (**Figure S1B**), fold-change values (fc) are randomly sampled from a gamma distribution:

$$\text{fc} \sim 1 + \Gamma(r_1, r_2)$$

where the shape parameter $r_1$ is drawn from a uniform distribution between 0.05 and 0.5 ($\text{Unif}(0.05, 0.5)$) and the scale parameter $r_2$ from $\text{Unif}(0, 0.1)$. This approach ensures that most fold changes are close to 1.0, generating a small proportion of DEHs while maintaining a majority of non-DEHs. Fold changes are then applied to either the control or treatment condition based on a random selection using a beta distributed sampling:

$$s_{\text{fc}} \sim \text{B}(2, r_3)$$

where parameter $r_3$ is sampled from $\text{Unif}(2, 6)$. If $s_{\text{fc}}$ <0.5, the fc is multiplied to $\mu^C$; otherwise, it is multiplied to $\mu^T$. This ensured that simulating DEHs with biased expression favoring one condition, for instance, more homeologs highly expressed under treatment conditions compared to control. Here, let $\boldsymbol{\mu}'^C$ and $\boldsymbol{\mu}'^T$ denote the mean of homeolog expression under control and treatment conditions after adjusting by fc.

At the same time, the homeolog expression ratio (HER) for each subgenome is generated with the following processes (**Figure S1C**). $N$ of $\theta_k$ are sampled from a normal distribution with the mean of $\frac{1}{K}$ and the standard deviation of $\left(\frac{1}{K}\right)^2$ and used for representing the baseline of the HER of subgenome $k$ for $N$ homeologs under control ($\boldsymbol{\theta}_k^C = \boldsymbol{\theta}_k$) and treatment ($\boldsymbol{\theta}_k^T = \boldsymbol{\theta}_k$) conditions. Then, to introduce HER variability, shifts in HER between conditions are simulated by drawing $\Delta\theta_k$ from a gamma distribution $\Delta\theta_k \sim \Gamma(1, 0.1)$ and rescaled into the range $\left[0, \frac{1}{K}\right]$. Next, $\Delta\theta_k$ are randomly added to $\theta_k^C$ or $\theta_k^T$ to mimic shifts in HER between control and treatment conditions. Let $\theta'^C_k$ and $\theta'^T_k$ denote $\theta_k^C$ and $\theta_k^T$ after adjusting with $\Delta\theta_k$. The sampling and rescaling ensure that most $\Delta\theta_k$ are close to 0, resulting in similar $\theta'^C_k$ and $\theta'^T_k$ for most homeologs pairs (**Figure S4** and **S5**). Once the processes are repeated from $k = 1$ to $k = K$, $\theta'^C_1, \ldots, \theta'^C_K$ and $\theta'^T_1, \ldots, \theta'^T_K$ are obtained. The values $\theta'^C_1, \ldots, \theta'^C_K$ are then normalized to sum to 1.0, and also for $\theta'^T_1, \ldots, \theta'^T_K$. Here, let $\boldsymbol{\theta}'^C_1, \ldots, \boldsymbol{\theta}'^C_K$ and $\boldsymbol{\theta}'^T_1, \ldots, \boldsymbol{\theta}'^T_K$ donate the simulated HER of $N$ homeologs of $K$ subgenomes under conditions C and T.

Finally, $\boldsymbol{\theta}'^C_k$ $(k = 1, \ldots, K)$ are multiplied to $\boldsymbol{\mu}'^C$ and $\boldsymbol{\theta}'^T_k$ $(k = 1, \ldots, K)$ are multiplied to $\boldsymbol{\mu}'^T$ to simulate the homeologs with different HER between conditions (**Figure S1D**). Then, small proportion of $\boldsymbol{\theta}'^C_k\boldsymbol{\mu}'^C$ and $\boldsymbol{\theta}'^T_k\boldsymbol{\mu}'^T$ was randomly changed to zero, inducing the suppressed expression of homeologs. The artificial read counts of subgenome $k$ under control and treatment conditions are finally sampled from NB distributions of $\mathcal{NB}\left(\boldsymbol{\theta}'^C_k\boldsymbol{\mu}'^C, \boldsymbol{\phi}_k^C\right)$ and $\mathcal{NB}\left(\boldsymbol{\theta}'^T_k\boldsymbol{\mu}'^T, \boldsymbol{\phi}_k^T\right))$, respectively (**Figure S1E**), where the dispersion $\boldsymbol{\phi}_k^c$ is calculated with the quadratic function $f$ by converting estimated variance to dispersion,

$$\boldsymbol{\phi}_k^c = \frac{(\boldsymbol{\theta}'^c_k\boldsymbol{\mu}'^c)^2}{e^{f(\log \boldsymbol{p}'^c_k\boldsymbol{\mu}'^c)} - \boldsymbol{\theta}'^c_k\boldsymbol{\mu}'^c}$$

To define ground truth RSH, $D_{max}$ and $OR_{max}$ were calculated from $\boldsymbol{\theta}'^C_k\boldsymbol{\mu}'^C$ and $\boldsymbol{\theta}'^T_k\boldsymbol{\mu}'^T$. $D_{max}$ is defined as the maximum absolute differences in HER between conditions for each subgenome:

$$\mathrm{D_{max}} = \max\left\{\left|\boldsymbol{\theta}'^T_1\boldsymbol{\mu}'^T - \boldsymbol{\theta}'^C_1\boldsymbol{\mu}'^C\right|, \ldots, \left|\boldsymbol{\theta}'^T_K\boldsymbol{\mu}'^T - \boldsymbol{\theta}'^C_K\boldsymbol{\mu}'^C\right|\right\}$$

Similarly, ORmax is defined as the maximum odds ratio among any pair of conditions in at least one subgenome. Then, ground truth RSH is defined as homeologs with $D_{max} > 0.2$ or $OR_{max} > 2.0$.
